# Zika Virus Infection of Murine and Human Neutrophils and their Function as Trojan Horses to the Placenta

**DOI:** 10.1101/2021.09.14.460378

**Authors:** NG Zanluqui, LG Oliveira, CM Polonio, TT França, GP De Souza, SP Muraro, MR Amorim, VC Carregari, C Brandão-Teles, Patrick da Silva, MG de Oliveira, RFO França, MP Cunha, ML Nogueira, D Martins-de-Souza, A Condino-Neto, JL Proença-Modena, JPS Peron

**Author notes:** Correspondence to: Peron JPS. Av. Prof Lineu Prestes, 1730 Lab 232 ICB IV Cidade Universitária, São Paulo -SP - Brazil. CEP 05508-000.; Modena JLP. Rua Bertrand Russel, s/n. Caixa Postal 6109 Campinas, São Paulo – SP – Brazil. CEP 13083-970. These authors contributed equally to this work (LG Oliveira, CM Polonio).

## Abstract

ZIKV is a 11Kb positive stranded flavivirus transmitted by infected *Aedes aegypti* and by sexual intercourse. After a short period of viremia of 5-7 days, the virus is cleared, and infection resolved in 80% of individuals. However, around 27% of the fetuses from pregnant infected mothers may develop several fetal brain and ocular pathology. Here we show that murine and peripheral blood human neutrophils support ZIKV infection and replication both *in vitro* and *in vivo*, which may correlate to the facilitation of vertical transmission. ZIKV did not interfere with cell viability, neither induced ROS production nor the release of NETs by infected neutrophils. Also, ZIKV infection of neutrophils did not trigger a pro-inflammatory profile, as evidenced by qPCR and proteomic analysis. Interestingly, ZIKV-infected neutrophils were isolated from the placenta were highly infected. The transference of *in vitro* ZIKV-infected neutrophils to pregnant female mice favored the transference of viral particles to the fetus. Conversely, neutrophil depletion with monoclonal antibodies reduced fetal viral loads whereas the treatment with recombinant G-CSF has the opposite effect. In summary, although it has already been shown that circulating monocytes harbor ZIKV, to our knowledge, this is the first report demonstrating the role of neutrophils during ZIKV infection, and most important, that it may act as a trojan horse to placental tissue directly impacting the pathogenesis of congenital syndrome.

## INTRODUCTION

Zika virus (ZIKV) is a 11Kb positive-stranded RNA enveloped flavivirus transmitted by the *Aedes egypti*(*1*) mosquito bite. In 2015, an outbreak started in Brazil causing soon spread to more than 90 countries worldwide causing a serious public health crisis leading the WHO to declare state of emergency of international concern(*2*). ZIKV is able to cross both placental and fetal blood brain barriers, reaching fetal brain tissue and causing significant death and early differentiation of neuronal precursor cells (NPCs)(*3–6*). Clinically, the ZIKV congenital syndrome (ZCS) is characterized by microcephaly, cortical atrophy, brain calcification, ventriculomegaly, taquigyria, associated or not to arthrogyposis and retinal, as reviewed(*7–8*). Although much has been discovered about the biology of the virus in the human host, still some gaps need to be fulfilled, specially concerning the mechanisms that orchestrate resistance or susceptibility to infection and neuropathology.

Neutrophils are the most abundant and one of the most important cells of the innate immune response, comprising around 50-70% of human peripheral blood cells(*9*). Neutrophils are involved in many diseases, from sepsis to cancer, autoimmunity and also viral infections, as Chikungunya(*10*) and SARS-CoV2(*11*). It has phagocytic and lytic capacity especially against extracellular bacteria and it may recognize many different pathogens, either by Toll-like receptors (TLRs) and RIG-like receptors (RLRs)(*12*). Intracellular killing mechanisms involve proteolytic enzymes, antimicrobial peptides, reactive oxygen species (ROS) and reactive nitrogen intermediates (RNI), as reviewed(*9*). It is recruited from the bone marrow after differentiation by GM-CSF and is capable of easily infiltrate tissues due to the expression of many chemokine receptors, mainly CXCR2 and its ligand IL-8/CXCL8, which plays a key role in neutrophilic chemotaxis during infections(*13*). Neutrophils may also undergo active cell death followed by chromatin decondensation and the release of DNA extracellular traps that actively destroy and trap pathogens(*14*). This phenomenon is called NETosis and may be both dependent and independent of ROS production(*15*). Also, as the first to invade damaged tissue, neutrophils actively secrete chemokines to recruit more inflammatory cells(*16*).

The relevance of neutrophils during viral infections have recently gained attention, as they play important roles during infection with viruses of the families Flaviviridae, Togaviridae, Orthomyxoviridae and Retroviridae(*16–20*). More interesting, they may either act on viral clearance (*18, 21, 22*), or as a reservoir for virus replication and dissemination (*18–20*). It has been demonstrated increased neutrophil counts during varicella zoster, Herpes simplex virus (HSV)(*23*), West Nile virus (WNV)(*18*) and influenza (PR8H1N1) virus infections(*22*). Corroborating this, the treatment with anti-Ly6G depleting antibody resulted in increasing viral loads in lungs and nasal fluid compared to controls of H1N1 infected mice(*24*). Also, neutrophil alpha-defensin may directly impair HSV-1/2 replication(*25*). Respiratory Syncytial Virus (RSV) induces the release of extracellular DNA traps that causes obstruction of the airways(*26*). This evidences the complex role of neutrophils in containing viral replication but also contributing for the pathogenesis of the infections.

On the other hand, some viruses may subvert neutrophils effector mechanisms, favoring viral persistence. For instance, human neutrophils are induced to live longer during cytomegalovirus (HCMV) infection. In fact, CMV genome codes for vCXCL-1, an analog of CXCL1 that recruits neutrophils to the site of infection and favor viral dissemination and pathogenesis (*27*). Concerning flaviviruses, Dengue virus (DENV) can also infect and replicate in neutrophils(*28*), whereas the brain pathology caused by WNV correlated with neutrophil hijacking(*18*). The role of neutrophils during ZIKV virus infection and ZCS has not been elucidated.

Here we show that both murine and human neutrophils are target for ZIKV replication *in vitro* and *in vivo* without interfering in their viability. More interesting, ZIKV infection did not induce ROS production neither the release of NETs. We also demonstrate that infected neutrophils may seed ZIKV to the placental and other tissues, favoring vertical transmission and pathology in mice. Conversely, this is corroborated after ZIKV infection in the presence of *Aedes egypti* saliva, a known recruitment factor for inflammatory cells to the bite site(*29*). Altogether, our data evidence the importance of neutrophils as possible viral factories during ZIKV infection that may act as trojan horses, transporting the virus to other organs, as brain and placenta. To our knowledge, this is the first report demonstrating the importance of neutrophils for ZIKV infection and vertical transmission.

## RESULTS

### ZIKV Infects and Replicates in Murine Neutrophils

To address whether ZIKV infects neutrophils we started by isolating bone marrow neutrophils from C57BL/6 and SJL mice and infected *in vitro* with ZIKV Brazilian strain BeH815744. Besides the anti-E protein, positive (gRNA) and negative (agRNA) ZIKV RNA genome were detected in neutrophils of both strains C57BL/6 (Fig 1A) and SJL (Supplementary Fig 2A) after 20 hours of infection. Corroborating this, ZIKV replication in neutrophils occurs as evidenced by PFU detection on the supernatant of ZIKV-infected neutrophils cultured for 48 hours (Fig 1B and Supplementary Fig 2B). Peak replication occurred 6 hours post infection (hpi) and progressively decreased until the end of the experiment. Consistently, ZIKV RNA was also detected in the cell lysates at 6 and 18 hpi (Fig 1C). The decrease in ZIKV particles releasing over the time is concomitant to the progressive cell death observed (Fig 1D and Supplementary Fig 2C). Interestingly, ZIKV infection did not change the viability of the cells compared to control (Fig 1D and Supplementary Fig 2C). Next, we addressed whether ZIKV would impair the neutrophil capacity of ROS production. As expected, neutrophils activated with PMA highly produced ROS, whereas ZIKV infected cells had ROS production at levels similar to the controls for 6 hours period in luminol oxidation assay (Figure 1E). This was corroborated by the decreased DCFDA^+^ cells from both C57BL/6 (Figure 1F) and SJL (Supplementary Fig 2D). In fact, ZIKV infection caused a slight decrease in the capacity of neutrophils to produce ROS in response to PMA, although only in the frequency of positive cells (Fig 1F). Additionally, observed that ZIKV decreased the phagocytic capacity of neutrophils evidenced by reduced Dextran-FITC uptake (Fig 1G). Next, to confirm the overall reduced neutrophilic activation during ZIKV infection, we evaluated the expression of genes related to host antiviral response, cell metabolism, neutrophil activation, degranulation and flavivirus cell attachment and invasion at 6 and 18hpi (Fig 1H and Supplementary Fig 2E and 3). Consistently, ZIKV did not change the expression of most of the genes analyzed, with reduction only of *Lcn2* (Fig 1H and Supplementary Fig 2E) and slightly of *Glut1* expression (Fig 1H and Supplementary Fig 3C) in C57BL/6 neutrophils. After 18 hours, ZIKV infection increased the expression of *Ifnb1*, *Nos2*, *Tnf-α* and *Axl* with the recovery of *Lcn2* and *Glut1* expression (Fig 1H and Supplementary Fig 3). Unexpectedly, the expression of most genes analyzed in SJL neutrophils after 9 hours of infection did not change besides an increase of *Ifnar2* (Supplementary Fig 2E). Further, as ZIKV is extremely susceptible to type I IFNs, and infected C57BL/6 neutrophils showed increased expression of *Ifnb1* without changing *Ifnar2* mRNA, we sought to evaluate the protein expression of type I IFN receptor 1 (IFNAR1) as well as the expression of other activation markers by flow cytometry. Confirming PCR finding, despite the increased expression of *Ifnb1*, *in vitro* infected-neutrophils showed no difference in the expression of CD80, CD86, MHC-I and II and IFNAR1 (Supplementary Fig 4).

**Figure 1:**
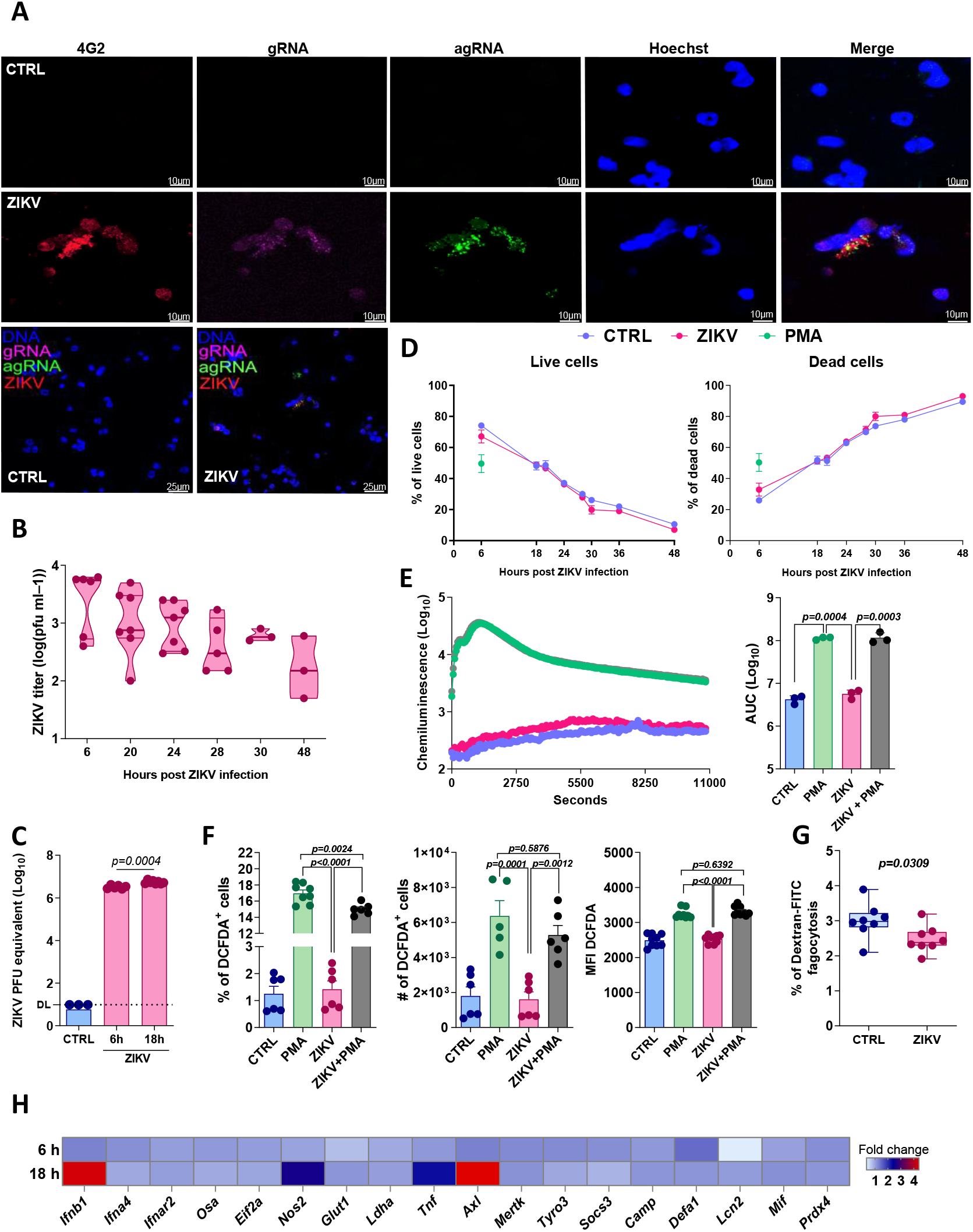
Zika virus infects murine neutrophils. A) Bone marrow neutrophils from C57Bl/6 mice were infected with ZIKV (MOI 1) for 20 hours and further submitted to prime flow staining for ZIKV genomic (gRNA - purple) and anti-genomic (agRNA - green) RNA detection and ZIKV envelope (ZIKV - red) staining with 4G2 and analyzed in confocal microscope. B) Plaque forming units (PFU) from ZIKV-infected neutrophils (MOI 0.1) cultures supernatants at the indicated time points. Representative of two experiments with at least three replicates each. C) At 6 and 18 hpi neutrophils were recovery and submitted to ZIKV RT-qPCR quantification (DL – Detection limit). Graph representative of three experiments with at least three replicates. D) Neutrophils were infected with ZIKV during the indicated time points or incubated with PMA for 30 minutes. Cell death was evaluated by Annexin V and 7-AAD staining by flow cytometry. Graph representative of three experiments with at least three replicates in each group. E) Neutrophil ROS production was evaluated in real time for 3 hours by oxidation of luminol and chemiluminescence in the presence of ZIKV and/or PMA. Graph of ROS quantification over the time (left) and Area Under Curve (AUC - right) of ROS quantification. Graph representative of three experiments with three replicates in each group. F) Neutrophil ROS production evaluated by flow cytometry with DCFDA after 6 hours of infection with ZIKV or PMA (last 30 minutes of incubation). In all analysis PMA (50nM) was used as a positive control. Data representative of three experiments with three replicates each group. G) Neutrophils were infected with ZIKV MOI 0.1 for 1h and then incubated with Dextran-FITC for 2.5h. Cells were washed and phagocytosis analyzed by flow cytometry. Graph representative of two experiments with four replicates each. H) Bone marrow neutrophils from C57Bl/6 mice were infected with ZIKV (MOI 0.1) for 6 and 18 hours and submitted to qPCR. Heatmap shows fold change of all genes normalized to Actb and control group. Graph representative of three experiments with at least three replicates. Unpaired two-tailed t-test was used in C and G. One-way ANOVA was used in E and F. Two-way ANOVA was used in H.

### ZIKV Infects and Replicates in Human Neutrophils and Does Not Induce NETs Release

To further corroborate our findings in murine neutrophils we used human peripheral blood neutrophils from healthy donors. Consistently, ZIKV E-protein, gRNA and agRNA ZIKV genome were detected in human neutrophils (Fig 2A). We confirmed that by the detection of endosomal viral factory of ZIKV particles inside neutrophils by electron microscopy (Fig 2B black arrow). Consistent to the results with murine cells, ZIKV infection of human neutrophils also released infective viral particles (Fig 2C) and significantly reduced their phagocytic capacity (Fig 2D). Consistently, a slight increase of *IFNAR2* and *NOS2* was observed in ZIKV-infected neutrophils 9 hpi, whereas no differential expression in genes related to neutrophil activation and degranulation, as *AZU1, DEFA1, LCB2* was detected (Supplementary Fig 5).

**Figure 2:**
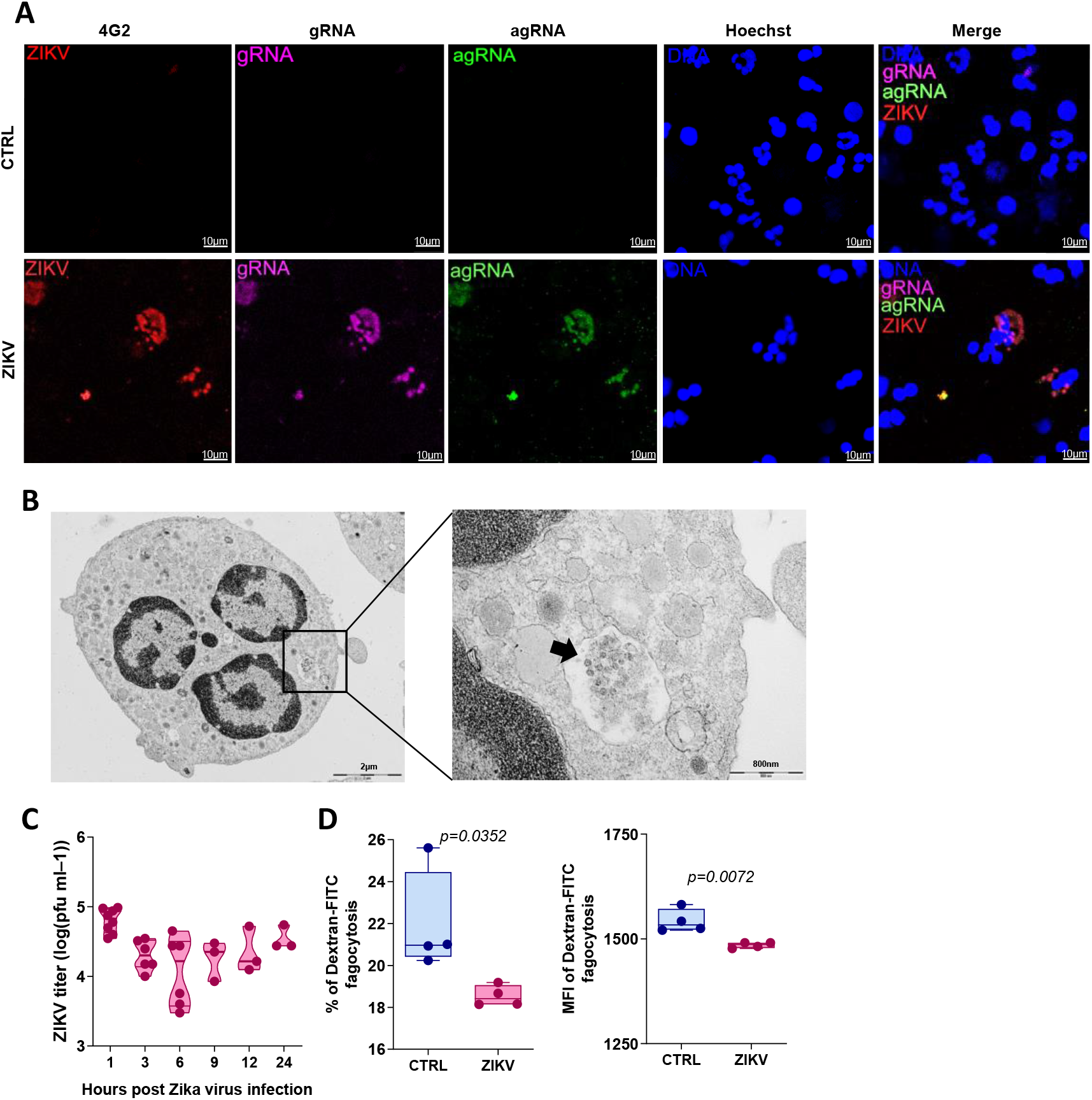
Zika virus infects human neutrophils. A) Peripheral blood neutrophils were isolated and infected with ZIKV (MOI 1) for 20h and submitted to prime flow staining for ZIKV genomic (gRNA - purple) and anti-genomic (agRNA - green) RNA detection and ZIKV envelope (ZIKV - red) staining with 4G2. The cells were analyzed confocal microscope, 63x of magnificence and 3 times zoom. B) ZIKV-infected neutrophils were submitted to electron microscopy. Images were acquired in 8900x (left) and 39Kx (right) of magnificence. C) Plaque forming units (PFU) detection of supernatants from ZIKV-infected neutrophils (MOI 0.1) at the indicated time points. Graph representative of three experiments with at least three replicates. D) Neutrophils were infected with ZIKV for 1h, then incubated with Dextran-FITC for 2.5 hours. After incubation, the cells were washed and analyzed by flow cytometry. Graph representative of two experiments with at least three replicates. Unpaired two-tailed t-test was used.

Next, we evaluated whether ZIKV was able to induce NETs release by neutrophil elastase (NE), histone H2A/H2B and DNA staining. After PMA exposure, histone H2A/H2B and DNA and NE could be detected spread in the cultures instead confined into the nucleus and granules, respectively (Fig 3A and B). Interestingly, ZIKV infection did not induce NETs release, evidenced by no signal of extracellular histone H2A/H2B and the intact structure of NE granules and nucleus similarly to control group (Fig 3A and B). Next, aiming to evaluate if ZIKV actively blocks NETs release, neutrophils were exposed to heat inactivated ZIKV (iZIKV) for 4 hours. No NETs were detected (Fig 3A). We also checked the ability of infected cells to release NETs after PMA stimulation in the presence of ZIKV or iZIKV exposure. The release of NETs after PMA stimulation in ZIKV+PMA, iZIKV+PMA and PMA alone was quite similar, despite a slight reduction of histone H2A/H2B in ZIKV+PMA and iZIKV+PMA (Fig 3A and B). We also confirmed that infected human neutrophils did not induce ROS production assayed by luminol oxidation for 3 hours (Fig 3C). Altogether, our results confirm that ZIKV infects and replicates in neutrophils without interfering with cell viability and without the induction of NETs release, although phagocytic activity was slightly decreased.

**Figure 3:**
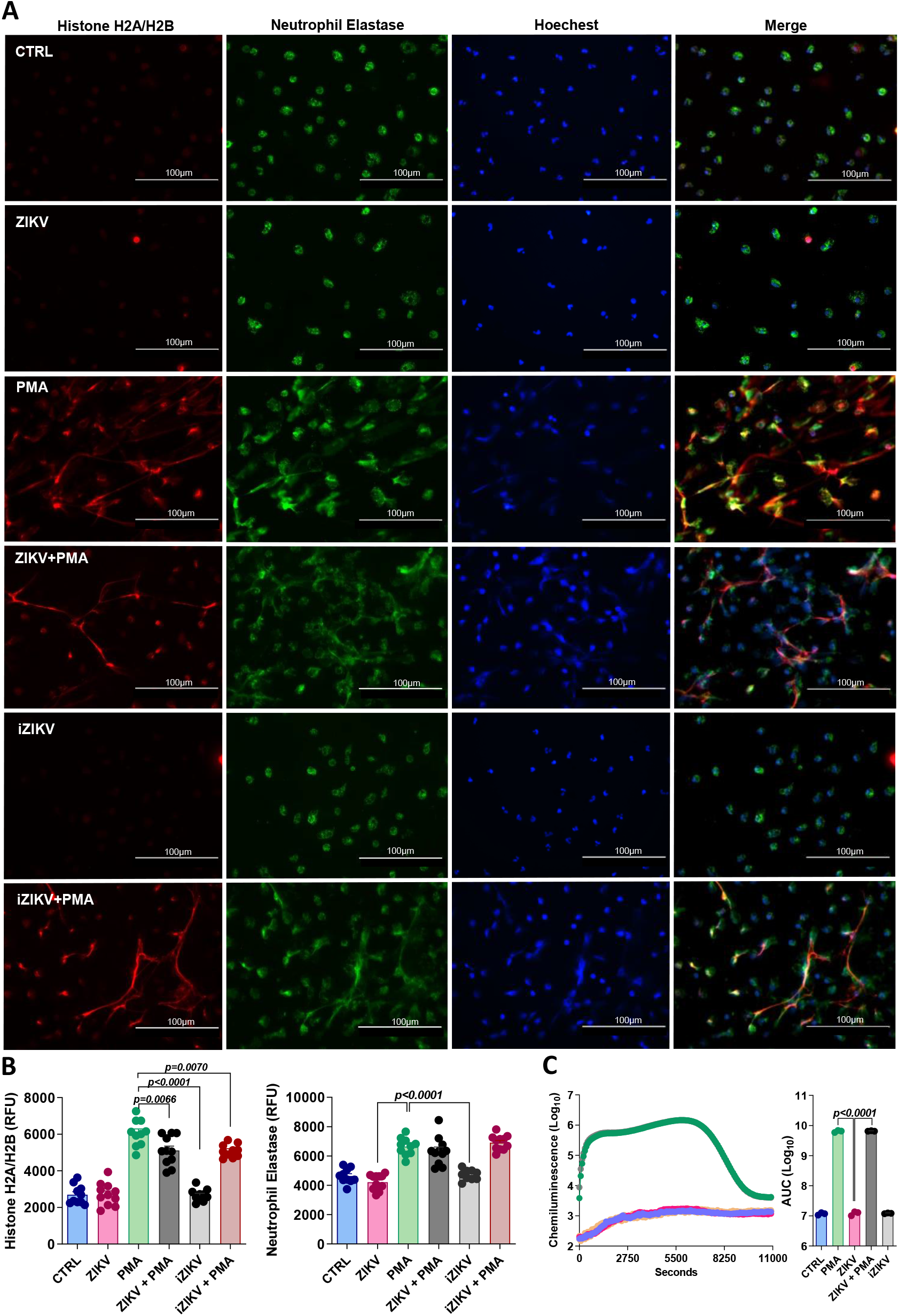
Zika virus infects human neutrophils and does not induce NETs release. A) Peripheral blood neutrophils were seedded in coverslips and infected with ZIKV or the same concentration of heat inactivated ZIKV (iZIKV) for 4 hours. After incubation, the cells were submitted to histone H2A/H2B (Orange), neutrophil-elastase (green) and Hoechst (blue) staining and analyzed by immunofluorescence. Pictures were acquired in 40x magnificence. B) Image quantification of histone H2A/H2B (left) and neutrophil-elastase (right) by ImageJ. n= 10 – 12 / group. C) Peripheral blood neutrophil ROS production was evaluated in real for 3 hours by oxidation of luminol and chemiluminescence in the presence of ZIKV, iZIKV and/or PMA. Graphic of ROS quantification over the time (left) and Area Under Curve (AUC - right). PMA (50mM) was used as positive control. Data representative of three independent experiments and from three different healthy subjects. One-way ANOVA was used in B e C.

### Neutrophils are important for ZIKV infection *in vivo*

Next, we evaluated whether neutrophils are relevant during ZIKV infection *in vivo*. For that, we infected C57BL/6 IFNAR1^-/-^ mice with ZIKV and 24 hours later peripheral blood neutrophils were isolated for ZIKV qPCR. Corroborating our data *in vitro*, E-protein, gRNA and agRNA were detected by prime flow (Fig 4A) and qPCR (Fig 4B). As expected, we confirmed that PBMCs were also infected, as previously shown (*45*) (Fig 4B).To further confirm the biological relevance of neutrophils during ZIKV infection *in vivo*, we treated IFNAR1^-/-^ mice with anti-Gr1 or anti-Ly6G antibody to deplete monocytes + neutrophils and neutrophils, respectively, prior to infection with ZIKV. As expected, twenty-four hours later, lower viremia was detected in mice depleted of neutrophils and monocytes (Fig 4C), as well as in neutrophils-depleted mice (Fig 4D), although in lesser extension. To further confirm this finding, IFNAR1^-/-^ mice were treated with rHu G-CSF (Filgrastim^®^) to increase the number of circulating neutrophils. In fact, increased neutrophil counts correlated with higher viral titters compared to controls (Fig 4E).

**Figure 4:**
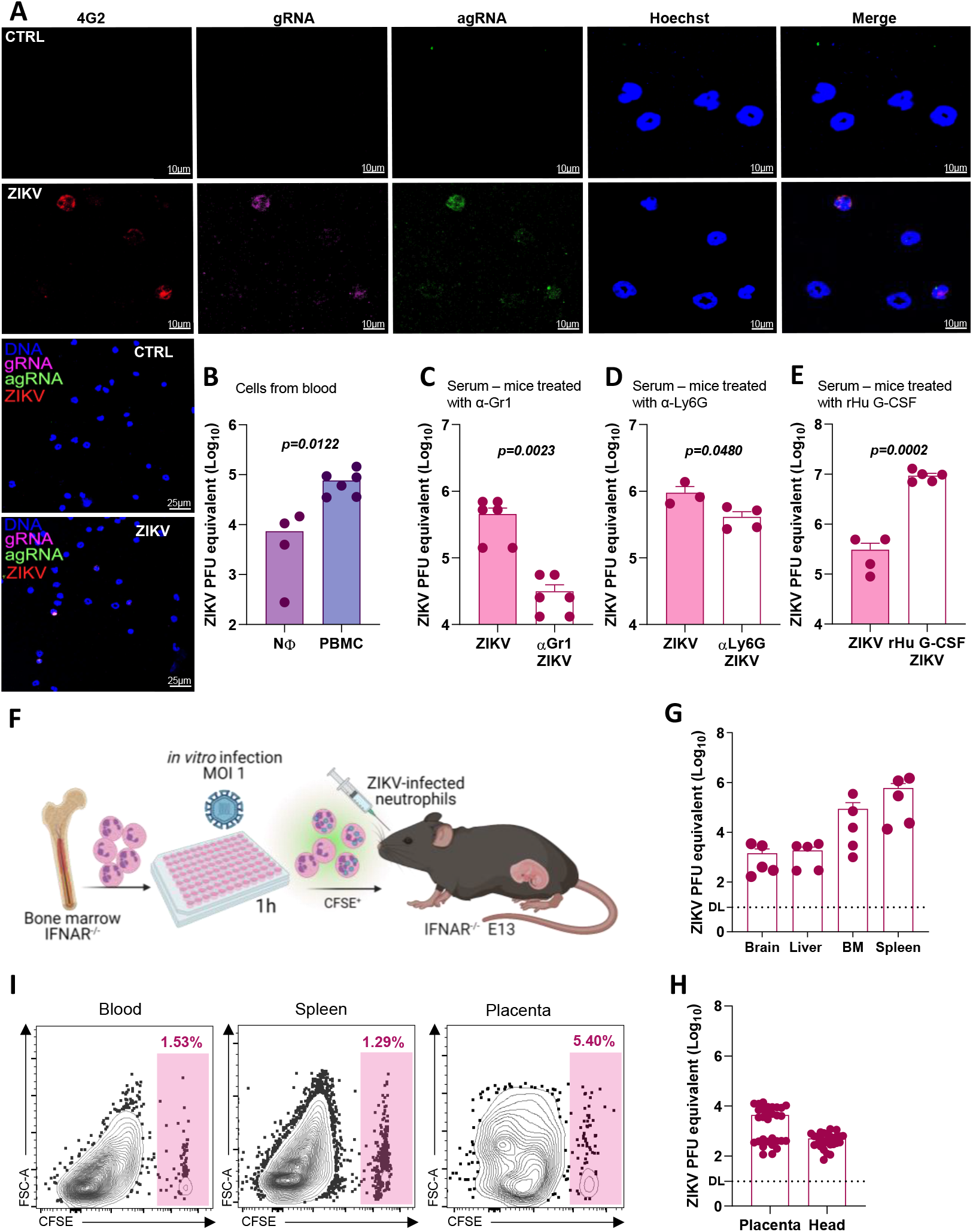
Neutrophils are infected in vivo and play key role during infection. A) IFNAR-/- mice were i.v. infected with 106 PFU of ZIKV and 24 hours after infection peripheral blood neutrophils were collected and submitted to prime flow staining protocol for ZIKV genomic (g) and anti-genomic (ag) RNA detection and ZIKV envelope (ZIKV - red) staining with 4G2. Cells were analyzed by a confocal microscope, 63x of magnificence and 3 times zoom. B) Neutrophils and polymorphonuclear (PBMC) cells were harvested from the blood and viral RNA quantified by RT-qPCR. Data representative of two experiments with at least 3 replicates each. C) IFNAR-/- mice were treated with anti-Gr1 antibody for 24 hours before infection. 24 hours later serum was collected, and viral RNA was quantified by RT-qPCR. Graph representative of two experiments. n = 6 mice/group. D) IFNAR-/- mice were treated with anti-Ly6G antibody 24 hours. 24 hours later serum was collected, and viral RNA was quantified by RT-qPCR. n = 3 - 4 mice/ group. E) IFNAR-/- mice were treated with human recombinant G-CSF 48 hours before infection. 24 hours later serum was collected, and viral RNA was detected by RT-qPCR. Graph representative of three experiments. n = 6 mice/group n = 4 - 5 / group. F) Experimental design. Bone marrow neutrophils from IFNAR-/- mice were harvested and infected in vitro with ZIKV MOI 1 for 1 hour. Cells were washed, stained with CFSE, and transferred to pregnant IFNAR-/- on gestational day 13 (E13). 24 hours later maternal and fetal tissues were harvested. Viral RNA detected by RT-qPCR in the G) brain, liver, bone marrow and spleen from the mothers and H) in the placenta and head of the fetus. Data from two experiments with 2 or 3 pregnant mice each. I) Blood, spleen, and placentas were harvested from F and cell suspensions were analyzed by flow cytometer for CFSE detection. Representative dot plot. Unpaired two-tailed t-test was used in B,C, D and E.

The recruitment of immune cells and the microinflammatory niche caused by the mosquito bite and saliva inoculation may improve viral infection, replication and dissemination *in vivo*(*16*). Noteworthy the fact that neutrophils are the main cells recruited to the skin after mosquito bites(*46*). Thus, we sought to evaluate whether the saliva from the ZIKV vector *Aedes aegypti* would play a role in facilitating infection in mice. For that, adult IFNARI^-/-^ mice were infected in the footpad with ZIKV alone or in the presence of *Aedes aegypti* saliva (Supplementary Fig 6A). Corroborating our hypothesis, we observed increased viremia in the saliva group at 24 hpi although no differences were observed in the paw, spleen, brain, liver and cells from the blood (Supplementary Fig 6B-G).

### ZIKV-infected neutrophils may disseminate virus to the placenta

Next, we sought to evaluate whether infected neutrophils can seed viral particles to the placenta and thus, playing an important role on the pathogenesis of CZS. As depicted (Fig 5F), bone marrow neutrophils were isolated and infected *in vitro* for 1 hour. Then, non-internalized viral particles were washed out and the cells transferred intravenously to pregnant IFNARI^-/-^ mice at embryonic day E13. To dissect transferred from host neutrophils, cells were stained with Carboxyfluorescein succinimidyl ester (CFSE). Mother and fetus’s tissues were evaluated 24 hours later (E14). ZIKV was detected in all mother tissues analyzed, as brain, liver, bone marrow and spleen (Fig 4G). More interesting, ZIKV was abundantly detected in the placenta and head of the pups (Fig 4H). Moreover, we confirmed the presence of the transferred neutrophils in the blood, spleen, and placenta (Fig 4I). These results evidence that ZIKV-infected neutrophils are present in the circulation and the placenta and that they may facilitate the dissemination of viral particles to fetal tissues.

**Figure 5:**
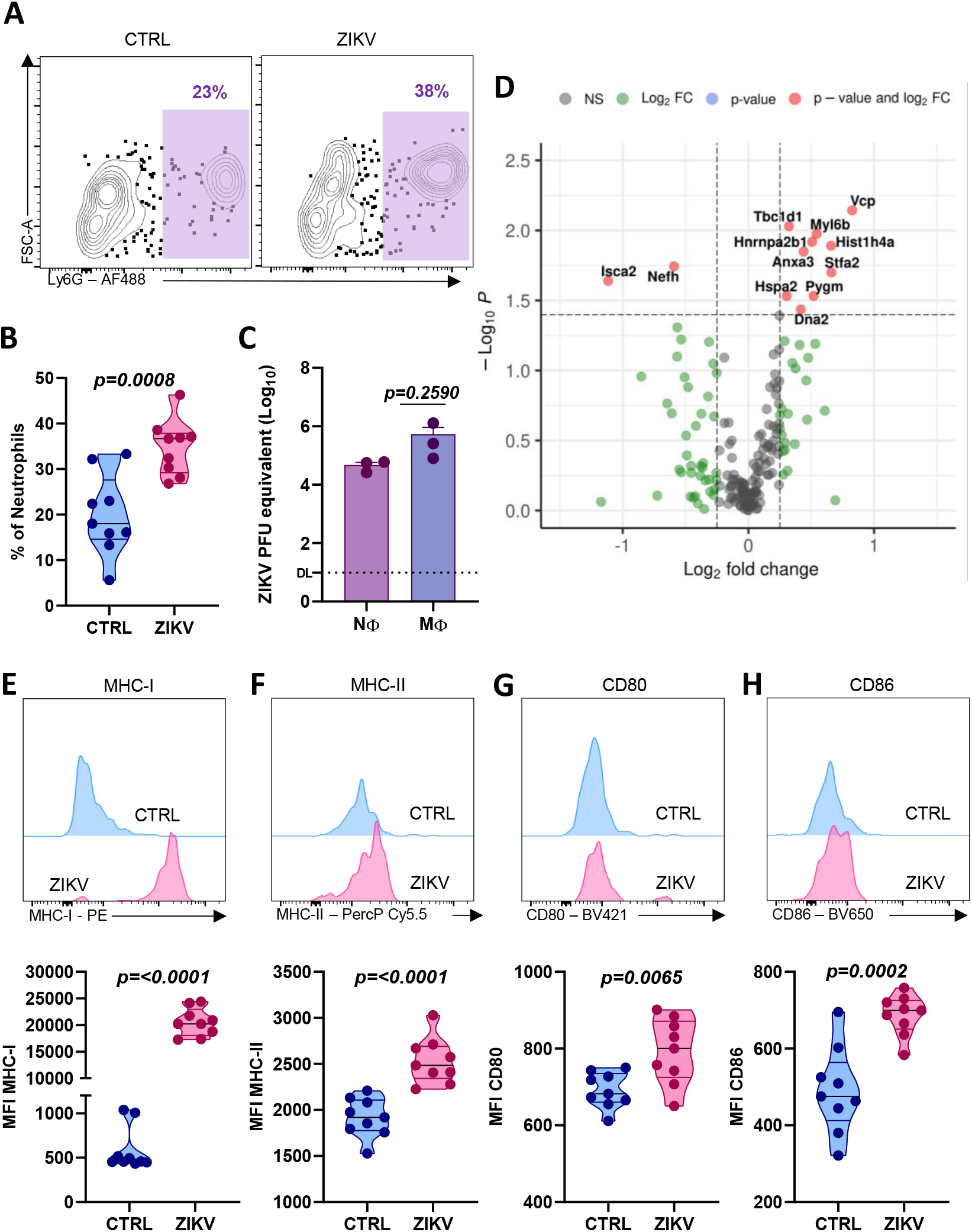
Placental neutrophils are infected with ZIKV and increase antigen presentation molecules expression. IFNAR-/- pregnant mice were infected at day E13 with 106 PFU of ZIKV and on day E17 placentas were harvested and dissociated for immunophenotyping and cell sorting. A) Representative dot plot and B) frequency of neutrophils from control (CTRL) and ZIKV-infected pregnant mice placenta. C, D)Placenta single cell suspension were submitted to cell sorting and neutrophils (gated on Ly6G+CD11b+) and macrophages (Ly6G-CD11b+) were submitted to C) mRNA extraction for viral RNA detection by RT-qPCR. D) Isolated neutrophils (gated on Ly6G+CD11b+) were submitted to protein analysis by proteomics. Volcano plot showing in orange 12 proteins differently expressed between control and ZIKV groups. E – H) Placental neutrophils gated on A and B were immunophenotyped. Representative histogram plot (on top) and the median of fluorescence intensity (MFI – on bottom) of E) MHC-I, F) MHC-II, G) CD80 and H) CD86. Unpaired two-tailed t-test was used in B, C, E to H.

### Murine placental neutrophils are infected but not activated

To confirm our previous data on the transference of neutrophils infected *in vitro*, we analyzed the frequency of neutrophils (CD11b^+^Ly6G^+^) and macrophages (CD11b^+^Ly6G^-^) in the placenta of IFNAR^-/-^ mice. As expected, both populations are present during all time points analyzed and their frequencies were similar during the time. However, whereas the peak of neutrophils frequency was at E18-19, absolute numbers peaked on day E16-17 (Supplementary Fig 7). This agreed with monocytes, although frequency decreased from E11-12 to E18-19 (Supplementary Fig 7).

Next, we analyzed the placentas of IFNAR1^-/-^ ZIKV-infected pregnant mice on E17 more profoundly. As expected, ZIKV infection significantly increased neutrophil frequency in the placenta (Fig 5A and B). Moreover, to address whether these cells were infected, we cell sorted neutrophils (CD11b^+^Ly6G^+^) and macrophages (CD11b^+^ Ly6G^-^) from the placentas and proceeded ZIKV RNA quantification. Consistent with previous findings, placental neutrophils harbor significant amount of virus, comparable to CD11b^+^Ly6G^+^ macrophages (Fig 6C). Differently from neutrophils infected *in vitro*, we also observed that placental neutrophils from ZIKV-infected mice had higher levels of antigen presentation markers as MHC-I (Fig 5E), MHC-II (Fig 5F), CD80 (Fig 5G) and CD86 (Fig 5H).

**Figure 6:**
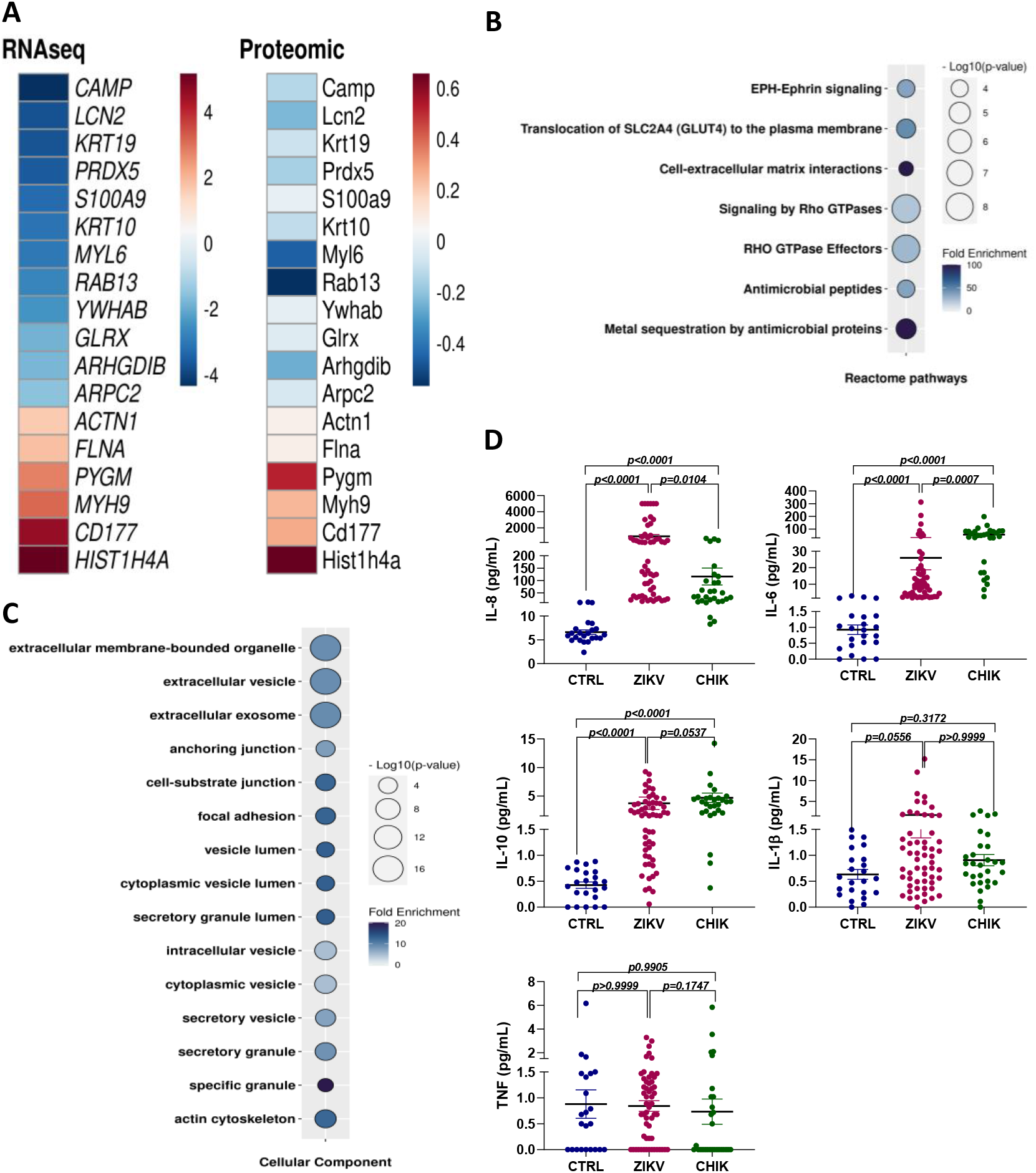
Comparison between mice placental neutrophils proteome and RNAseq of human placental tissue. Available RNAseq data set from Lum et al., 2019 of CD45+ cells from pre-term placenta of ZIKV-infected women in the first trimester and health donor was compared to our sorted placental neutrophil proteomics. A) heatmap with human genes (on left) and murine proteins (on right) that were concordant in both data sets. The first 12 were downregulated (in blue scale), whereas the last 6 were upregulated (in red scale). B) GO of reactome pathways and C) Cellular componentsof genes and proteins showed in A. D) Cytokines in the serum of ZIKV and CHIV-infected patients and healthy individuals. CTRL n = 23; ZIKV n = 58; CHIV n = 29. One-way ANOVA and Kruskal-Wallis multiple-comparisons test was used.

Considering that neutrophils from placenta were highly infected, we sought to evaluate their proteomic profile by sorting CD11b^+^Ly6G^+^ cells and submitting to proteomic analysis. Interestingly, of a total of 192 proteins identified (Supplementary Fig 8), only 12 were only slightly differentially expressed between control and ZIKV-infected placental neutrophils (*p<0,05*) (Fig 5D). 10 proteins were upregulated, whereas 2 were downregulated in ZIKV-infected neutrophils (Fig 5D). This seems consistent with our data of *in vitro* infected neutrophils evidencing an absence of a significant translational activation of these cells despite the increase in antigen presentation molecules.

### Human placenta datasets evidence enrichment of neutrophil related genes with downregulation of activation markers

As previously shown, around 80% of CD45^+^ cells in the ZIKV-infected human placenta are neutrophils (*47*). To correlate our findings in mice with human samples, we compared data sets of CD45^+^ cells from placentas of healthy and ZIKV-infected women in the first trimester of pregnancy. We focused on the expression of genes related to neutrophil function, activation, and degranulation. Corroborating our findings, we found that all these genes were significantly downregulated in ZIKV-infected placenta (Supplementary Fig 9A). Conversely, gene ontology (GO) analysis of downregulated genes in ZIKV-infected placenta showed enrichment of genes of immune response pathways, such as leukocyte, granulocyte and neutrophil activation and degranulation and cytokine-mediated signaling (Supplementary Fig 9B). Moreover, viral infection processes pathways such as viral gene expression e transcription were also enriched as downregulated genes (Supplementary Fig 9C). Altogether, these analyses showed that ZIKV infection promote downregulation of genes related to immune response and neutrophils function in the placenta, although also downregulates viral infections processes.

To further confirm this correlation, we compared our proteome analysis of murine placental neutrophils with the same RNAseq data set from the CD45^+^ cells of ZIKV infected human placentas (Supplementary Fig 9). In fact, we can observe that there is a significant convergence of molecules downregulated in both conditions, although with slight difference in the level of expression (Figure 6A). More interesting, GO analysis evidences the downregulation of several cellular activation processes, as cell-extracellular matrix interactions, signalling by Rho GTPases, Rho GTPases effectors and antimicrobial peptides (Figure 6B) as well as several secretory pathways, as secretory granule lumen, secretory vesicle, cytoplasmic vesicle, and many others. These data indicate that despite neutrophils are abundant and infected in placental tissues of ZIKV patients, they seem not to be activated, but rather, suppressed or modulated.

Finally, as IL-8/CXCL8 has a key role on neutrophils recruitment (*13*) and inflammatory cytokines are important during arbovirus infection, we sought to determinate cytokine levels in ZIKV patients and compared to CHIKV patients and control subjects. Using the inflammatory CBA kit we detected IL-1β, IL-6, IL-8, IL-10 and TNF-α. As expected, high levels of IL-8 and IL-6 were detected in both infections, although ZIKV-infected patients showed higher IL-8 levels. Interestingly, the opposite was observed for IL-6 (Fig 6D). IL-10 was increased in both infections. No differences were observed for IL-1β, IL-10 and TNF-α between ZIKV- and CHIKV-infected subjects (Fig 6D).

## DISCUSSSION

ZIKV as well as many other flaviviruses represent a great threat for human health. The outbreak of Zika virus started in Brazil in 2016 and put the world in state of concern, as declared by WHO on February 1st. The virus was soon correlated with a skyrocketing increase of babies born with microcephaly, as well as adults diagnosed with Guillain-Barré Syndrome(*7*) and other neurological features(*48*). Currently, Brazilian public health system supports around 3200 microcephaly babies, in addition to many more with visual or auditive impairments. Although the crisis has ended, it is indispensable that scientific research continues for a better understanding of the pathogenesis of CZS and the development of therapeutic interventions.

Here we show that neutrophils are important cells for the biology of ZIKV infection, and possibly for the pathogenesis of the CZS. Although the function of neutrophils against fungi and bacterial infections is well stablished (*49, 50*), their role during viral infections has been neglected for a long period, but has recently gained attention (*11, 19, 51*). The antiviral activity of neutrophils is controversial, as it may be beneficial to the host, as in herpes simplex(*23*), Marburg and Ebola virus(*52*) infections, or detrimental, as in CHICKV (*10*), WNV(*18*), Yellow Fever Virus (YFV) and even SARS-CoV2(*11*). The main reason is because neutrophils may be hijacked by viruses, supporting their replication and dissemination(*18–20*), or because their effector mechanisms when exacerbated may lead to tissue damage (*10, 11, 27*). This evidences the complexity of neutrophil function during viral infection and the need for further investigations.

Mosquito’s saliva is composed of several molecules, including NeSt1 protein (neutrophil-stimulating factor 1) that potently stimulates IL-1β and CCL2 secretion, with further neutrophilic infiltration to the site of infection, promoting viral replication and increasing disease severity(*16,46,53*). Local immune inflammation recruits myeloid cells, including neutrophils and macrophages, that may act as virus reservoir and carriers to the lymph nodes and other tissues. Corroborating this, we observed that ZIKV infection in the presence of *Aedes egypti* saliva significantly increased viremia in mice. Conversely, depletion of circulating neutrophils with monoclonal antibodies anti-Gr1 and anti-Ly6G significantly reduced the viral load, whereas the enrichment of the neutrophil population after the treatment with G-CSF had the opposite effect. In agreement with that, we observed high levels of IL-8/CXCL8, a well-known chemokine for neutrophil recruitment, in the serum of ZIKV patients when compared to controls or CHIKV-infected patients.

One important difference between the biology of CHIKV and ZIKV infection is that CHIKV patients develop severe arthritic responses, with swollen joints and acute pain, whereas these symptoms are rarely observed during ZIKV infection. Arthritogenic mechanisms rely on the intense release of NETs by neutrophils and low density neutrophils that actively participate not only in the pathogenesis of CHIKV infection but also on rheumatic diseases, as systemic erythematous lupus (SLE)(*54*) and juvenile idiopathic arthritis (JIA) (*55*). Here we show that neutrophils participate differently in the pathogenesis of ZIKV infection, i.e., not through the activation of effector mechanisms, as NETs release and ROS production, but rather by acting as a trojan horse to the placental tissue.

Here we suggest that, in addition to CD16^+^ monocytes(*45*), neutrophils also exert important function for ZIKV dissemination *in vivo*. It is noteworthy that the frequency of neutrophils in the blood of humans and mice is very different. While in human blood neutrophils comprise about 50-70% of leukocytes, in mice this percentage drops to 10-25%(*56*). Despite these differences, neutrophils are target of ZIKV infection in both cases which may indicate that the importance of neutrophils for human infection is even higher. By using anti-Gr1 antibody prior to infection, depleting both neutrophils and monocytes (*57*), we observed a drastic reduction in viremia, which shows us that both cell populations are relevant for viral replication. To test neutrophils specifically, we used anti-Ly6G antibody, and we also observed a reduction in viremia, although in lesser extent. Furthermore, pretreatment with human recombinant G-CSF had an opposite effect, increasing viremia.

Interestingly, DENV hemorrhagic patients express high levels of CD66b, a marker for cell activation and NETs formation, associated with high IL-8/CXCL8 levels. (*28*). Conversely, neutrophils from milder disease patients do not produce NETs(*28*). It has been shown that *in vitro* infection of neutrophils with DENV-2 interferes with NETs formation by reducing the expression of Glut-1 and, consequently, the uptake of glucose, which is important for NETs release(*17*). This agrees with our findings, as we also observed that ZIKV infection reduced ROS production and NETs release by neutrophils, associated with slightly reduction of Glut1 mRNA and many other activation markers. It was also demonstrated that HIV envelope glycoprotein induces high IL-10 levels by dendritic cells which is capable of inhibiting NETs formation and ROS production (*21, 58*). Accordingly, we observed that despite higher IL-8/CXCL8 plasma concentration, ZIKV-infected individuals significantly higher circulating IL-10 than healthy individuals, which was also observed in infected women(*59*). However, IL-10 levels were similar between ZIKV and CHIKV patients, indicating that other mechanisms may be present to suppress NETs release. Also, interferon stimulated genes (ISGs) *Osa, Eif2* were downregulated although the expression of *Ifnb1* was increased. No difference was detected in the protein expression of the IFN receptor in *in vitro*-infected neutrophils, and neither in peritoneal neutrophils from *in vivo*-infected mice (data not showed), which may significate that post-transcriptional control may take place.

Furthermore, *in vitro* infection with ZIKV does not induce the expression of cell activation markers, including MHC-I. However, peritoneal (data not showed) and placenta neutrophils from infected mice have high levels of CD80, CD86, MHC-I and MHC-II, evidencing the influence of the microenvironment on neutrophil activation. Worth to mention that our findings agree with the literature, as a high frequency of neutrophils were detected in the placentas of ZIKV infected women. Besides, our proteomic analysis when compared to published data sets confirm that there is absence of neutrophil activation in the placenta, which may support the idea of neutrophils as trojan horses.

One weakness of our study is that we were not able to sample human ZIKV infected placentas to evaluate neutrophils. However, we compared the proteome of our murine sorted neutrophils from placental tissue with data sets from CD45^+^ cells obtained from from human placenta of ZIKV infected patients. Interestingly, there was a great concordance on the enrichment of genes related to neutrophils, but, most importantly, the expression of genes related to neutrophil activation, degranulation and interaction with extracellular matrix were downregulated. This may indicate that placenta neutrophils are not only inactivated, but they may have been hijacked by ZIKV. In fact, it has already been demonstrated that leukocyte count during pregnancy is high(*60*) as well as that healthy pregnancies correlate with reduced phagocytic and oxidative functions of neutrophils, probably in order to maintain maintaining immunosuppression and fetal acceptance (*61*). In mice, the recruitment of neutrophils to the placenta correlate with a better pregnancy outcome due to the local induction of iTregs. Neutrophil depletion with anti-Ly6G during pregnancy not only reduced decidual CD4^+^Foxp3^+^ Tregs, but also had a high impact on placental and fetal sizes(*62*). Conversely, it was demonstrated that neutrophils comprised up to 80% of CD45^+^ cells in the placenta of ZIKV-infected individuals, which is approximately 20% more than the healthy placenta(*47*). This may be explained by the high expression of IL-8/CXCL8 in the fetal side of the ZIKV-infected placenta, promoting the active recruitment of neutrophils. Corroborating this, we showed that the transference of *in vitro*-infected neutrophils to pregnant mouse promotes ZIKV vertical transmission, as we detected viruses not only in the placenta, but also in the head of the fetuses. Although it has already been well described that ZIKV infects macrophages, trophoblasts and Hofbauer cells of placental chorionic villi (*63, 64*), to our knowledge, this the first time that ZIKV were detected in placental neutrophils. In fact, our findings agree with a recent description of placental histopathology from ZIKV infected women evidencing moderate to intense neutrophilic chorioamnionitis that correlated with higher levels of calcifications and bleeding(*65*). Noteworthy that placental bleeding and damage mediated by type I interferons has been implicated in the facilitation of vertical transmission and fetal pathology in mice (*66*). Whether type I interferons derive from placental neutrophils, although plausible, still needs to be determined.

Altogether, our research brings attention to neutrophils concerning the pathogenesis of ZIKV infection, and especially for CZS. To our knowledge, their role as a trojan horse to the placental tissue is also a novelty. In this sense, we believe our data contributes for a better understanding on the pathogenesis of ZIKV infection, mostly to the mechanisms of susceptibility to fetal pathology, and posing neutrophils as important players that must be further investigated.

## MATERIALS AND METHODS

### Murine bone marrow neutrophil isolation

Neutrophils were isolated from the hind paws bone marrow of C57BL/6, SJL and IFNAR^-/-^ mice. Red blood cells were removed by hypotonic lysis and to the cell pellet was added 2mL mL of Percoll^®^ 55%. The gradient was assembled with 5 mL of Percoll^®^ 81%, followed by 5 mL of Percoll^®^ 62% and finally, the cells were gently added to the top of the gradient. Cells were centrifuged at 24°C, 1200g and without brake for 30 minutes. After centrifugation, the neutrophils were collected and resuspended in RPMI-1640 medium without phenol red (LGC^®^ Biotechnology) with 3% fetal bovine serum (FBS) and 1% penicillin and streptomycin (pen/strept). Isolation yielded around 70 - 80% neutrophils. All animal procedures were approved by Ethics Committee in Animal Procedures of the ICB-USP - CEUA #4714050719.

### Human neutrophils isolation

Heparinized blood was obtained from healthy volunteers by venous puncture after informed consent. Neutrophils were isolated after 6% dextran sedimentation and Ficoll-Paque Plus (GE Healthcare^®^) density gradient separation. Cells were centrifuged at 24°C at 900g without brake for 20 minutes. Residual erythrocytes were removed by hypotonic lysis, and neutrophils were resuspended in RPMI-1640 without phenol red with 3% FBS and 1% pen/strept. Isolation yielded around 95 - 98% neutrophils. All procedures were approved by the Ethics Committee of the ICB-USP (#2.612.149).

### ZIKV stock preparation and neutrophil infection

Zika virus Brazilian isolate (BeH815744(*30*)) was propagated in C636 cells (ATCC® CRL-1660™) and tittered in Vero-E6 cells (ATCC® CRL-1586™) by plaque-forming unit assay (PFU). Isolated neutrophils were seeded in 24 or 96 wells plates and infected with different multiplicity of infection (MOI) of ZIKV for 1h at 37°C with 5% of CO2. In some experiments, ZIKV was heat inactivated at 60°C for 2 hours. After infection, the virus was washed out and RPMI-1640 without phenol red with 3% FBS and 1% pen/strept was added to the cells and incubated for different time points.

### Plaque forming units (PFU)

To quantify replicating viral particles, VERO-E6 cells (ATCC^®^ CRL-1586 ™) were seeded in 24 well plates with RPMI-1640 medium (LGC Biotechnology^®^) 2% FBS and 1% pen/strept. The samples were diluted and added to the culture for 1 h in a CO2 incubator at 37 ° C. Further, the medium removed and semisolid medium was added (2x DMEM high glucose (LGC Biotechnology^®^) with 3% Carboxymethylcellulose (CMC), 2% of SFB and 1% pen/strept and incubated at 5% CO2 and 37 °C. Five days later, the cells were fixed with 4% formaldehyde (LabSynth^®^) and stained with 1% Violet Crystal (LabSynth^®^) for plaque visualization. Viral titers were expressed as plaque forming units (PFU) per milliliter.

### Reactive Oxygen Species (ROS) detection

To detect ROS production, neutrophils were incubated with 10μM of DCFDA (DCFDA - Cellular ROS Detection Assay - Abcam^®^) for 30 minutes. Cells were then centrifuged, and the pellet resuspended in RPMI without phenol red with 3% FBS and 1% pen/strept. The assay was carried out under different conditions: medium alone, ZIKV (MOI 0.1), PMA (50ng/ml) and ZIKV+PMA. After 6h of incubation, the cells were washed and analyzed using Attune^®^ flow cytometer (Thermo Fisher^®^).

Neutrophil respiratory burst was also measured by monitoring the luminol oxidation by chemiluminescence, as previously described(*31*). Briefly, 10^5^ neutrophils/well were seeded in 96 wells plate and incubated under different conditions: medium alone, ZIKV (MOI 0.1), PMA (50ng/ml), ZIKV+PMA and iZIKV (MOI 0.1) in the presence of luminol (1 mmol) (Sigma-Aldrich^®^). The chemiluminescence was monitored for 3 hours using a microplate luminometer reader (EG&G Berthold^®^ LB96V, Bad Wildbad, Germany). The results were expressed as relative light units (RLU) and area under the curve (AUC).

### Cell death assay

Isolated neutrophils (5×10^5^ cells per well) were incubated in three different conditions: medium alone, ZIKV (MOI 0.1) or PMA (50ng/ml) in RPMI-1640 without phenol red with 3% SFB and 1% pen/strept in a CO2 incubator at 37°C for each timepoint. After incubation, the cells were washed with 1X Annexin buffer, and the cell pellet was resuspended in annexin buffer with Annexin V-FITC and 7-AAD according to the manufacturer’s specifications (FITC Annexin V 7-AAD-Thermo Fisher^®^). Flow cytometry was performed using Attune flow cytometer (Thermo Fisher^®^). All singlets were analyzed for Annexin V and the 7-AAD labeling. Analysis was performed using FlowJo^®^.

### Flow cytometry

After 18 hours of infection, neutrophils were incubated with anti-CD16/32 (clone 93 - FcBlock), MHC-I (clone 34-1-25), anti-MHC-II (clone M5/114.15-2), anti-Ly6G (clone 1A8), anti-CD80 (clone16-10A1), anti-CD86 (clone GL-1), anti-CD11b (clone M1/70) and anti-IFNAR1 (clone MAR1-5A3) in the presence of live and dead (Zombie red^TM^ Dye Biolegend) for 30 min at 4 °C. Next, cells were washed, fixed with paraformaldehyde 1% and acquired using a Fortessa LSR^®^ flow cytometer (BD Bioscience^®^). Doublets were excluded and the median of expression of CD80, CD86, MHC-I and MHC-II in CD11b^+^Ly6G^+^ gated cells were analyzed. Peritoneal macrophages from intraperitoneal ZIKV-infected mice were used as positive control. Analysis was performed in FlowJo^®^.

### Immunofluorescence

Immunofluorescence for the detection of NETs was performed as described by Brinkmann et al., 2010(*32*). Briefly, 2.5×10^5^ neutrophils/well were seeded in 13 mm diameter coverslips for 1 h in CO_2_ incubator at 37°C. Further, cells were infected with ZIKV MOI 0.1 and incubated for 1 h. Then, PMA (50 ng/mL) was added as positive control for NETs release. After 3 hours, cells were fixed with 4% paraformaldehyde. The permeabilization was performed with 0,5% Triton X-100 and blocked with 3% AB human serum, 1% BSA and 0.05% Tween 20 in PBS during 30 min at 37 °C. Cells were incubated with primary antibodies anti-histone H2A/H2B (antibody gently donated by Prof. Arturo Zychlinsky, Max Planck Institute, German) and anti-neutrophil elastase (Calbiochem^®^) for 1 hour in CO_2_ incubator at 37°C. Next, the cells were washed and incubated with secondary antibodies Alexa Fluor 488 and Alexa Fluor 546 (Invitrogen^®^) for 1 h in CO_2_ incubator at 37°C. Hoechst 33342 (Santa Cruz Biotec^®^) was added, and slides were assembled using Mowiol mounting medium and analyzed using Evos Cell Imaging System (Thermo Fisher^®^).

### Prime flow for Zika virus detection

Primeflow^®^ was carried out according to the manufacturer’s specifications and adapted for immunofluorescence. Briefly, ZIKV-infected neutrophils were incubated with fixation buffer 1 for 30 minutes and washed 3 times with permeabilization buffer. After that, the cells were stained with 4G2 antibody (anti-ZIKV E protein) followed by overnight incubation with fixation buffer 2. Next, cells were incubated with RNA Pre Amp mix and RNA Amp mix for 1 hour each. Label probes were added to the cells and incubated for 1 hour. Finally, cells were washed and Hoechst 33342 (Santa Cruz Biotec^®^) was added and the slides assembled using Mowiol mouting medium and analyzed by Leica TCS SP5 II confocal microscope.

### Phagocytosis assay

Neutrophils were infected with MOI 0.1 for 1 hour, washed and then incubated with Dextran-FITC (0.5mg/mL) for 2.5 hours. Next, cells were washed with PBS and acquired using Accuri^®^ C6 flow cytometer (BD Bioscience^®^) flow cytometry and analyzed using FlowJo^®^.

### Placenta dissociation and neutrophil isolation

Placenta dissociation was performed as described by Arenas-Hernandez et al., 2015(*33*). Briefly, placentas were collected in cold PBS 1X with 1% pen/strep. To prepare cell suspensions, placentas were chopped in 3 mL of Accutase solution (Thermo Fisher^®^) and incubated in 80RPM/min for 30 min at 37°C. Next, RPMI 10% FBS was added to the cells and passed through 70 µm mesh and centrifuged at 500g for 5min. Pellets were resuspended in 37% of Percoll with RPMI without phenol red, centrifuged at 1200g for 20min without break and resuspended with RPMI and submitted to cell sorting. Cell suspensions were stained with anti-CD11b-PE (clone M1/70) and anti-Ly6G Alexa-fluor 488 (1A8) in the presence of live and dead (Zombie red^TM^ Dye Biolegend) and anti-Fc receptor (CD16/32 – clone 2.4G2) for 30 minutes. Next, the cells were washed and resuspended in RPMI with 10% of SFB and EDTA 0,05mM and submitted to cell sorting in Moflo Astrios^®^ cytometer (Beckman Coulter^®^). The gate strategy and neutrophil yield are supplied in the supplementary figure 1.

### Transference of neutrophils to pregnant mice

Bone marrow neutrophils from female IFNAR^-/-^ mice were isolated using Percoll gradient and 2×10^6^ cells were infected with ZIKV MOI 1 for 1 hours. After infection, the virus was washed out and the cells resuspended in 100µL of PBS for intravenous adoptive transference to IFNAR^-/-^ female mice on the embryonic day (E)-13 of pregnancy. Twenty-four hours later, the animals were euthanized, and mother’s spleen and brain and pup’s head and placenta were collected for viral RNA detection by qPCR.

### G-CSF treatment and neutrophil depletion

To increase the number of circulating neutrophils, IFNAR^-/-^ mice of 6 and 10 weeks were intravenously treated with 140ug/kg of Filgrastim^®^ (rHu G-CSF – Blau Farmacêutica) diluted in glucose 5% solution. Two days later, the animals were infected with 10^6^ PFU of ZIKV and twenty-four hours later the blood was analyzed for viral RNA quantification by RT-qPCR. Neutrophil increase was monitored by hemogram. For the depletion of neutrophils, IFNAR^-/-^ mice between 6 and 10 weeks were intravenously treated with 200ug of antibody anti-Gr1 (clone RB6-8C5) or 50ug of anti-Ly6G (clone 1A8). One day later, animals were infected with 10^4^ PFU of ZIKV and twenty-four hours later, the blood was analyzed for viral RNA quantification by RT-qPCR.

### Infection with *Aedes aegypt* saliva

IFNAR^-/-^ mice between 6 and 10 weeks were infected subcutaneously with 10^6^ PFU of ZIKV in the presence or not of 10 µg of *Aedes aegypt* saliva. Twenty-four hours later, polymorphonuclear cells, the paw, brain, liver, and spleen were harvested for viral RNA detection by qPCR.

### Cytometry Beads Array

Serum from ZIKV- or CHIKV-infected and health donor were collected and submitted to cytokine detection by cytometric beads array using inflammatory human kit that detects IL-8, IL-1β, IL-6, IL-10, TNF and IL-12p70, following the manufacture’s specifications. Briefly, the serum was incubated with capture beads for 1.5 hours, washed and detection beads were added and incubated another 1.5 hours. Next, beads were washed and acquired in Accuri C6 (BD Biosciences^®^) and the analysis were performed in FCAP Array software (BD Biosciences^®^).

### Electron microscopy

Isolated neutrophils from health donor blood were infected in vitro with ZIKV MOI1 for 18 hours. Next, cells were recovered, washed with PBS 1x and fixed with Glutaraldehyde 4% overnight. Samples were washed with PBS 1x and followed to dehydration process with acetone gradient, desiccation, and gold metallization. Finally, sample of neutrophils were analyzed in JOEL1010 transmission electron microscope.

### mRNA extraction, cDNA, and q-PCR analysis

mRNA was extracted using RNeasy Mini kit^®^ (Qiagen^®^) or Arcturus^TM^ PicoPure^TM^ (Applied Biosystems^®^) following manufacturer’s specifications. RNA concentration was determined using NanoDrop (Thermo Fisher^®^) under 260/280nm wavelengths. High-Capacity cDNA Reverse Transcription Kit (Life Technologies^®^) was for cDNA synthesis. qPCR was performed using TaqMan^®^ gene expression assay (Thermo Fisher^®^) or SYBR Green^®^ (Thermo Fisher^®^) as described in the results.

TaqMan primers:

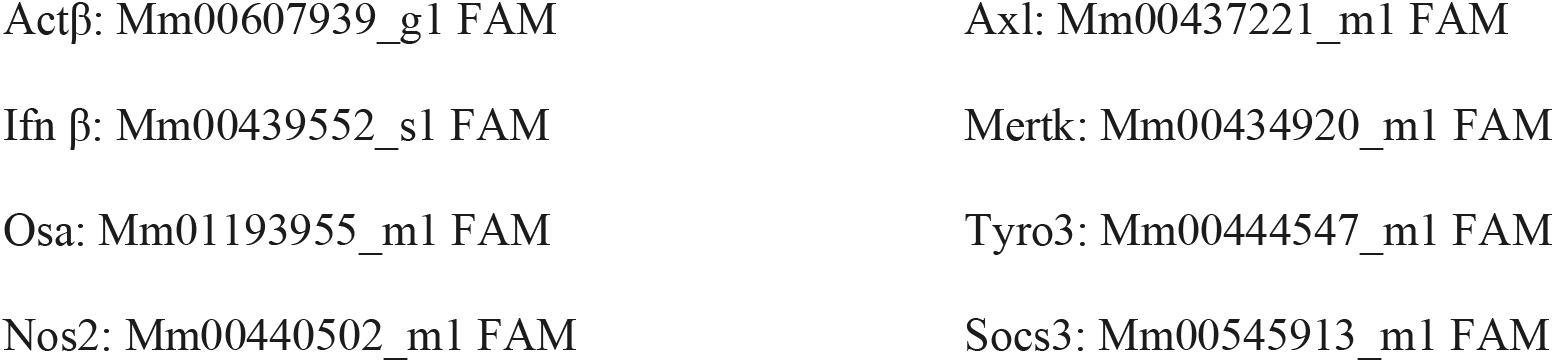
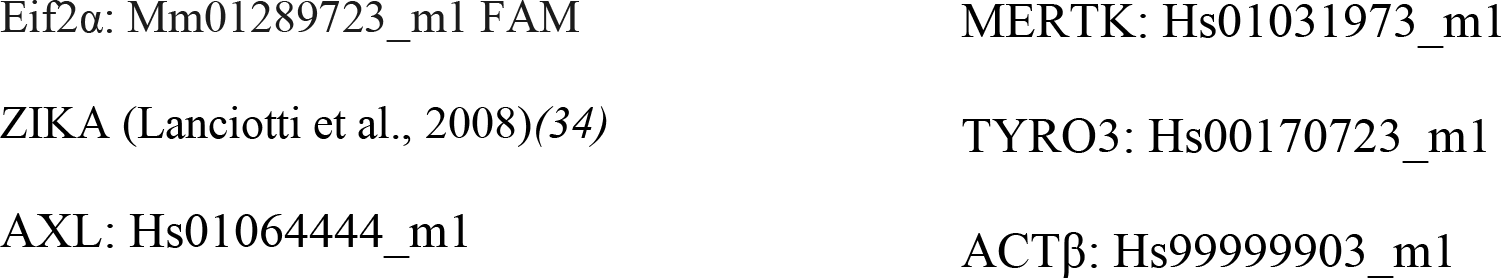

SYBR Green primers sequence:

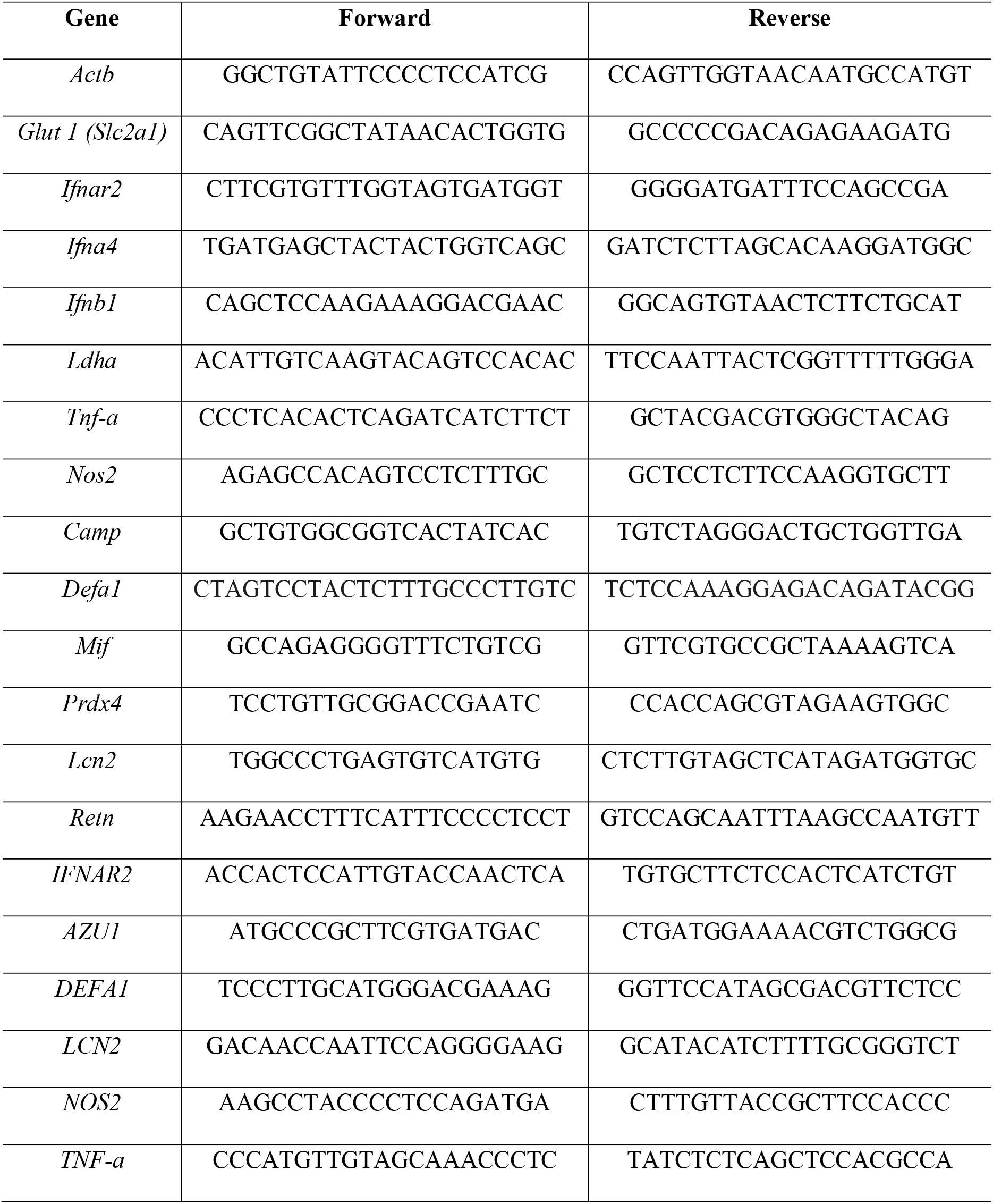

### Proteomics analysis

#### Sample preparation

After isolation, neutrophils from placenta were incubated with lysis buffer (100mM Tris-HCL pH 8, 150mM NaCl, 1mM EDTA, 0,5% triton) and washed. Proteins were digested with trypsin (Promega^®^) following filter-aided sample preparation (FASP) protein digestion as described by Wisniewski, JR et al., 2011(*35*). Briefly, to each filter sample YM-10 (Millipore^®^) was added urea (8M urea, 0,1 M Tris/HCl, pH 8,5) and the samples were span down twice. Then, proteins were reduced by 8mM (dithiothreitol) DTT and alkylated by 0.05M iodoacetamide (IAA) in urea (8M), and filters and samples were washed 3 times. Next, trypsin digestion was carried out at 37°C overnight in 50mM Ammonium bicarbonate (Ambic pH=8.0). The day after, the filters were transferred to a new tube and 50mM Ambic was added. Digestion was blocked by adding formic acid (FA) to a final concentration of 1% (v/v). The solutions were lyophilized and followed to the LC-MS analysis.

#### Mass spectrometry

Digested peptides have been analyzed by High Definition Synapt G2-Si (Waters Corporation^®^) mass spectrometer directly coupled to the chromatographic system (LC-MS). Differential protein expression was evaluated with a data-independent acquisition (DIA) of shotgun proteomics analysis by High-Definition Expression configuration mode (HDMSe). The mass spectrometer operated switching between low (4eV) and high (15-40eV) voltage collision energies on the fragmentation cells, using a scan time of 1.5s per function over 50-3000 m/z. All spectra were acquired in ionic mobility by applying a wave velocity for the ion separation of 1000 m/s and a transference wave velocity of 175 m/s. The processing of low and elevated energy added to the data reference mass peptide ([Glu1] -Fibrinopeptide B Standard, Waters Corp.) provide a time-aligned inventory of accurate mass retention time components for both the low and elevated-energy (EMRT, exact mass retention time). Search parameters were set as: automatic tolerance for precursors and product ions (20 ppm), minimum of 3 fragments combined per peptide, minimum 3 fragments pared per protein, minimum of 2 peptide combined per protein, 2 missed trypsin-cleavage, carbamidomethylation of cysteines and methionine oxidation as fixed and variables modification, false positive rate (FDR) of the identification algorithm below to 1%.

#### Label-Free Quantification of peptides

The analysis of differential proteins expression was done accordingly to Silva JC et al., 2006(*36*) and Visser et al., 2007(*37*). The data from LC-MS from 3 technical replicates for each sample were processed for qualitative and quantitative analysis in Progenesis QI software (Waters Corp). The expression analysis was performed considering technical replicated available for each experimental condition following the hypothesis that each group is an independent variable. The normalized protein table were generated to include protein identified at least in 2 of the 3 technical replicates and with switch regulation ± 20%; only proteins with p ≥0,05 were considered.

#### Mapping of RNA-seq libraries and differential gene expression analysis

RNA-seq raw data was obtained from the dataset GSE139181 on Gene Expression Omnibus public databank. For this study purposes, we selected and compared only CD45^+^ placenta discs data acquired from healthy and first trimester ZIKV infected pregnant women. Initially raw reads served as input for Trimmomatic(*38*), which performed quality filtering removing Illumina adaptor sequences, low quality bases (phred score quality > 20), and short reads. Trimming was followed by read error correction by SGA *k*-mer-based algorithm(*39*). Next the reads were mapped against the Genome Reference Consortium Human Build 38 (GRCh38) using HISAT2 software(*40*) following previously described optimization(*41*).The gene differential expression calling was achieved initially by counting the number of reads in each transcript through HTSeq(*42*). Finally, the count data were direct to differential analysis with DESeq2 R package(*43*). Biological process Gene Ontology terms search were performed through online database PANTHER with standard parameters(*44*).

### Statistical analyses

All statistical analyses were performed using GraphPad Prism software (Graphpad Software Incorporation^®^) version 6. The differences between the means of the results were determined by Student’s *t-test* and multiple comparisons were made by one-way ANOVA, depending on the number of variables. Significance level was set at p <0.05.

## AUTHOR CONTRIBUTIONS

Conceptualization, NGZ, JLPM and JPSP; Methodology, NGZ, LGO, CMP, TTF, SPM, MRA, GPS, VCC and MGO; Proteomics data analysis, VCC, CBT, PS and DMS; Patient enrolling and sample acquisition TTF, MPC and MLN; Original draft NGZ; Review and editing JLPM and JPSP; Funding acquisition JLPM and JPSP; Resource RFOF, ASN, DMS, ACN, JLPM and JPSP; Supervision JLPM and JPSP.

---

**Supplementary figure 1:**
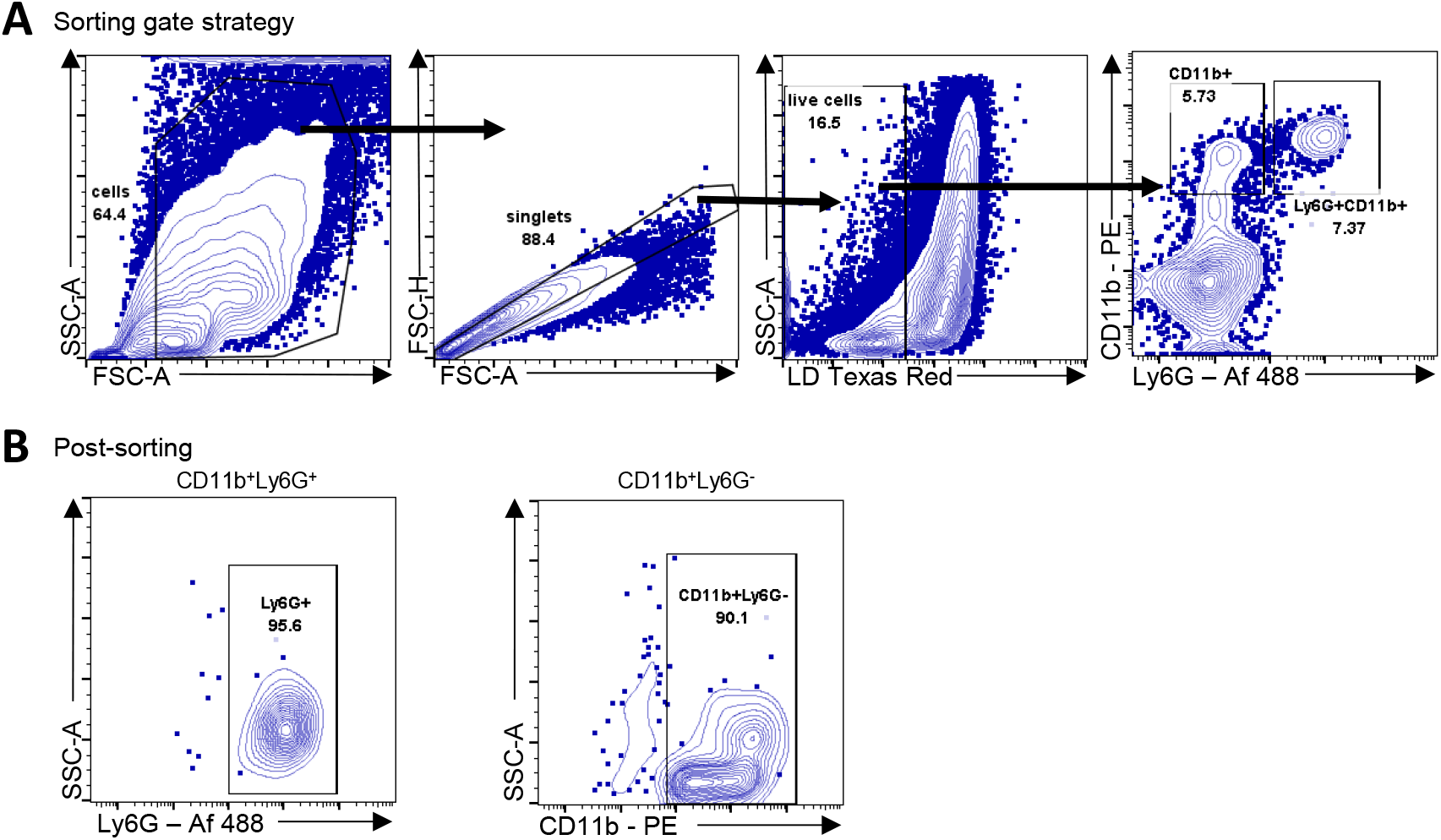
Neutrophil isolation from murine placenta. A) Gate strategy for placental neutrophil isolation by cell sorter. B) Neutrophils (CD11b+Ly6G+) and macrophages (CD11b+Ly6G-) cells isolation yield after cell sorter used in figure 5.

**Supplementary Figure 2:**
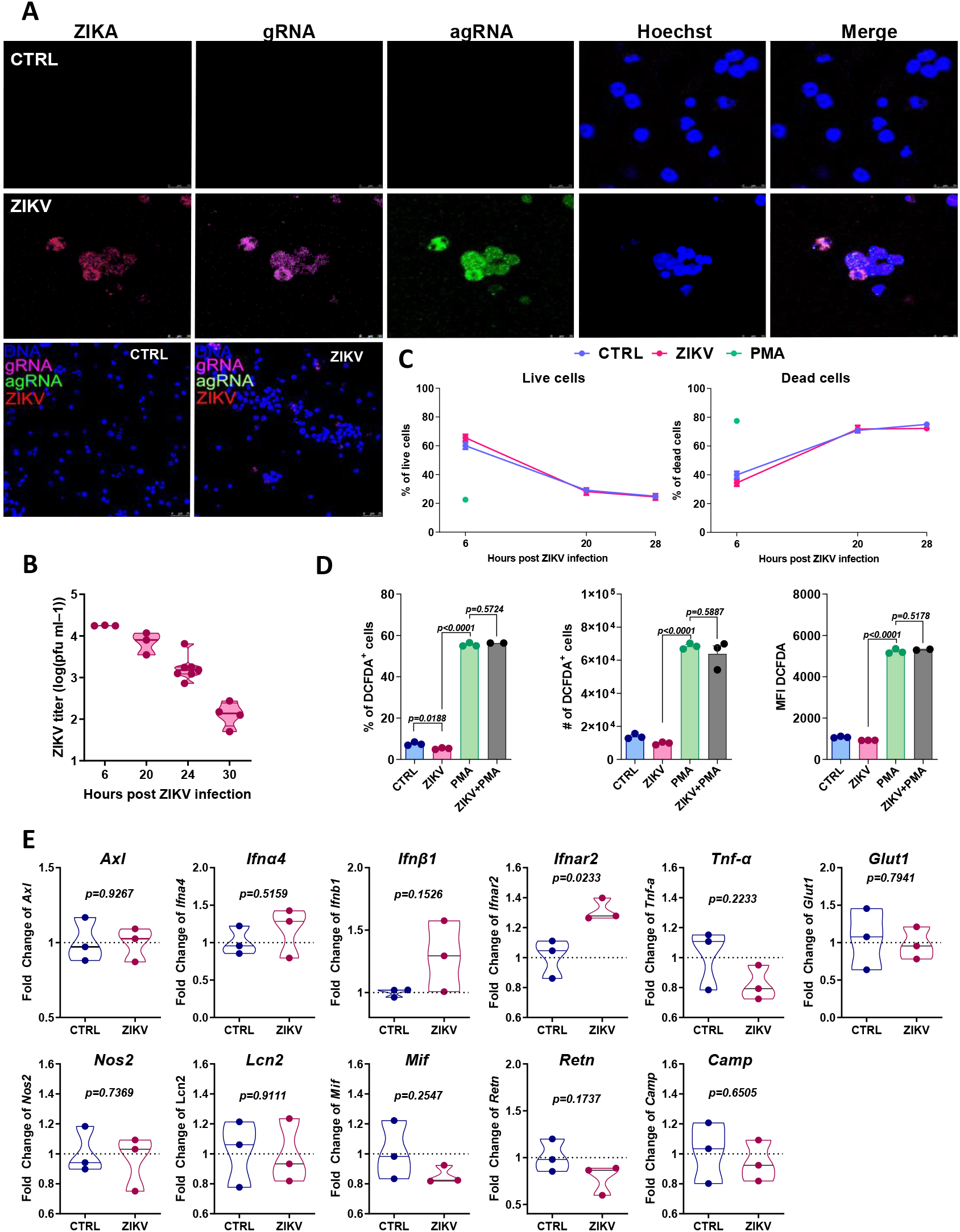
Zika virus infects SJL neutrophils. A) Bone marrow neutrophils from SJL mice were infected with ZIKV MOI 1 for 20 hours and further submitted to prime flow staining protocol for ZIKV genomic (gRNA - purple) and anti-genomic (agRNA - green) RNA detection and ZIKV envelope protein (ZIKV - red) staining. Cells were analyzed using confocal microscope. Pictures were acquired in 63x of magnificence and 3 times zoom. B) Supernatants from ZIKV-infected neutrophils cultures were collected at the indicated time points and submitted to PFU assay. Graph representative of two experiments with three replicates each. C) ZIKV-infected neutrophils were incubated during indicated time points and cell death analysis was performed by flow cytometry with Annexin V and 7-AAD staining. D) Neutrophil ROS production was evaluated by oxidation of DCFDA by flow cytometry. Neutrophils were incubated with DCFDA for 30 minutes at 37⁰C and further infected with ZIKV MOI 0.1 for 6h. During the last 30 minutes of infection PMA (50mM) was added to respective wells and used as positive control. Experiment repeated two times with three replicates each. E) Neutrophils were infected with ZIKV (MOI 0.1) for 9 hours and submitted to qPCR for Axl, Ifna4, Ifnb1, Ifnar2, Tnf, Glut1, Nos2, Lcn2, Mif, Retn and Camp genes. Fold change of all genes normalized by Actb and control group. One-way ANOVA was used D. Unpaired two-tailed t-test was used in E.

**Supplementary Figure 3:**
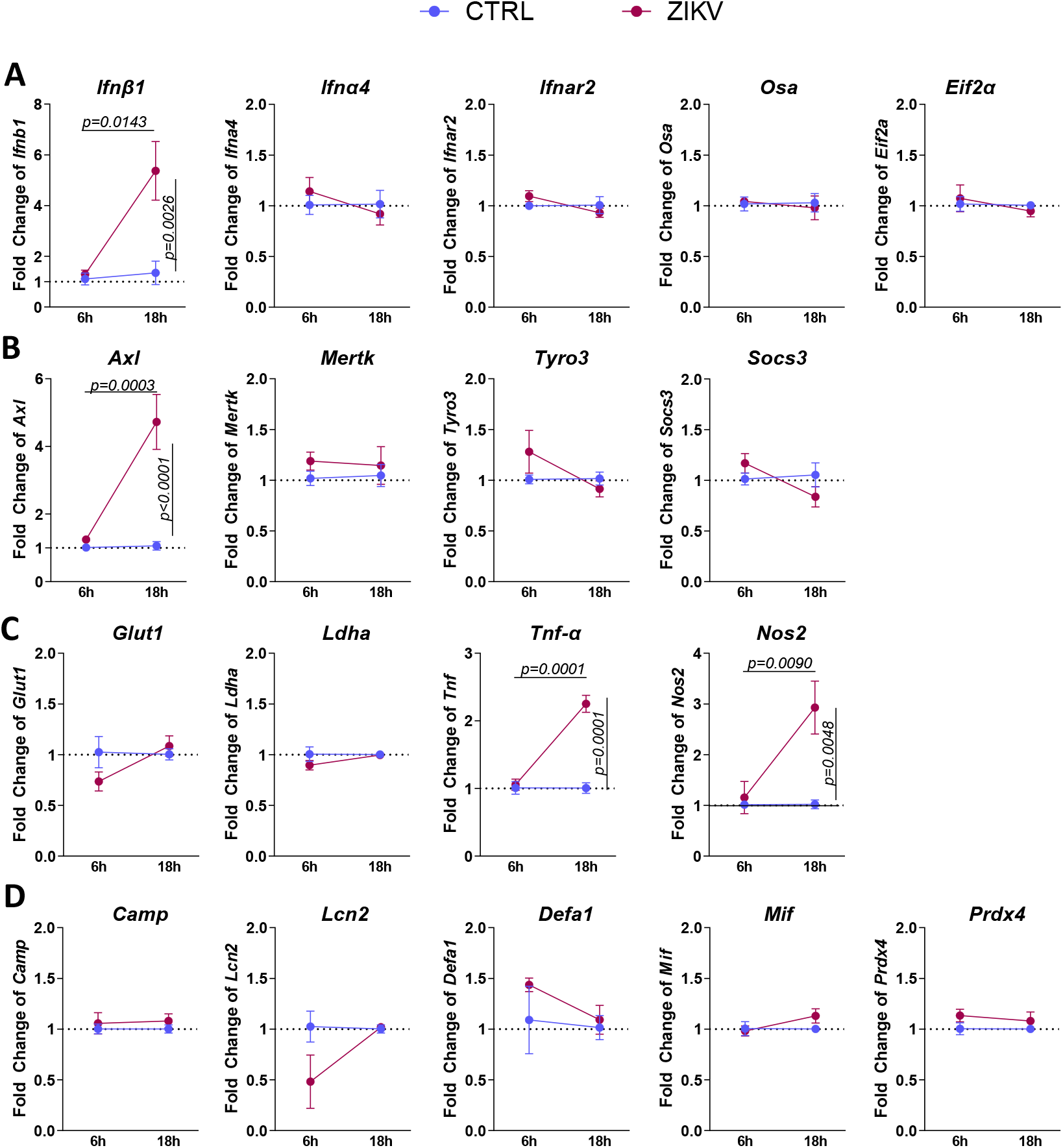
Gene expression profile in ZIKV-infected neutrophils. Bone marrow neutrophils from C57Bl/6 mice were infected with ZIKV (MOI 0.1) for 6 and 18 hours. Then, the total RNA was extracted and submitted to qPCR for A) Interferons and Interferon stimulated genes (Ifnb1, Ifna4, Ifnar2, Osa and Eif2a), B) TAM receptors (Axl, MertK, Tyro3 and Socs-3), C) Metabolism and immune response (Glut1, Ldha, Tnfa and Nos2) and D) Neutrophil function (Camp, Lcn2, Defa1, Mif and Prdx4). Fold change was normalized by Actb and control group. Data used in the heatmap showed in figure 1H. Data representative of 1-3 experiments with 3 replicates each. Two-way ANOVA.

**Supplementary Figure 4:**
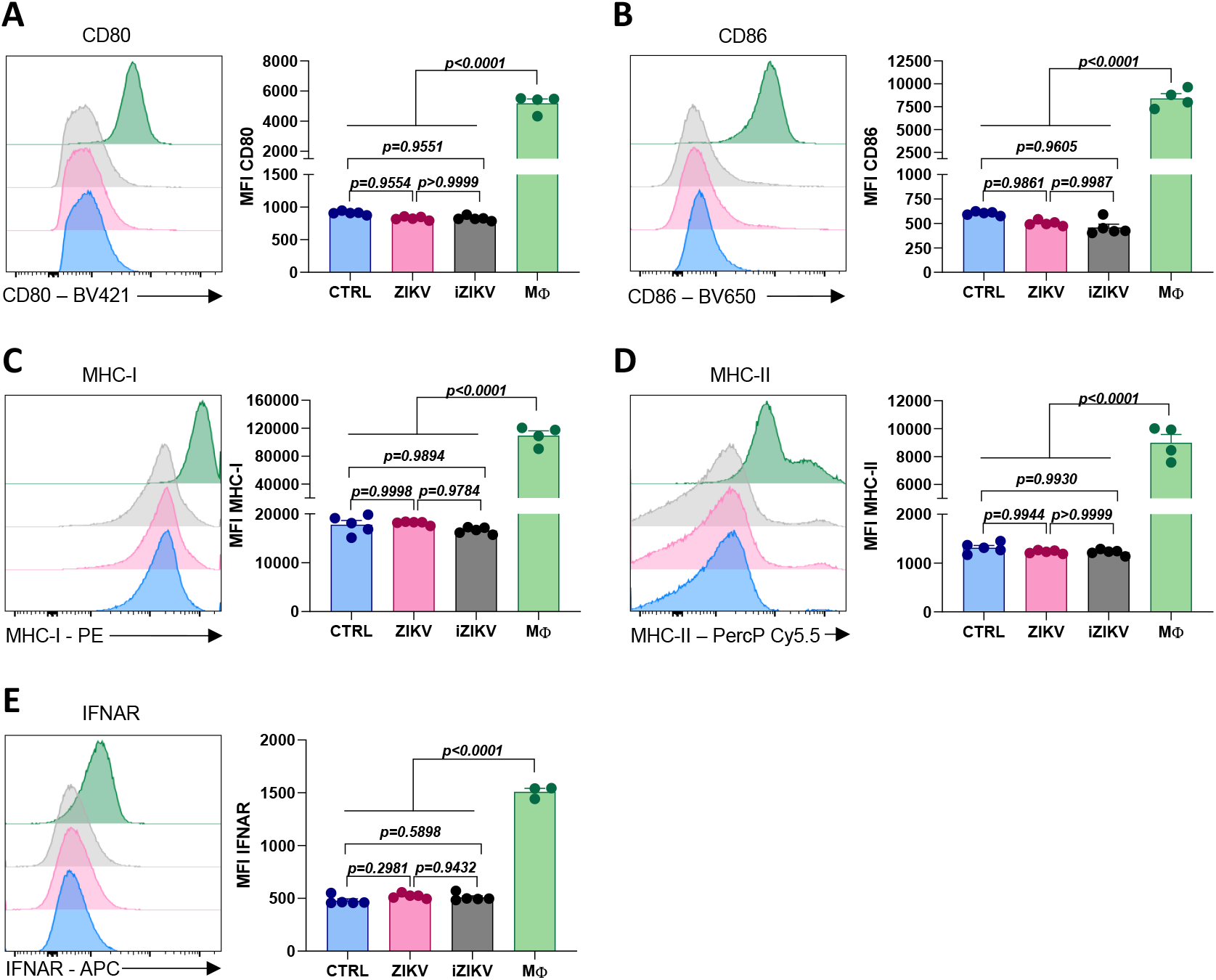
Antigen presentation molecules expressed by in vitro ZIKV-infected neutrophils. Bone marrow neutrophils were isolated from C57BL/6 mice and infected in vitro with ZIKV MOI 1 for 18 hours. After that, cells were recovered and stained for A) anti-CD80, B) anti-CD86, C) anti-MHC-I, D) anti-MHC-II and E) anti-IFNAR. Peritoneal macrophages from C57BL/6 i.p. Infected mice were used as positive control. Experiment with 3-5 replicates. One-way Anova was used.

**Supplementary figure 5:**
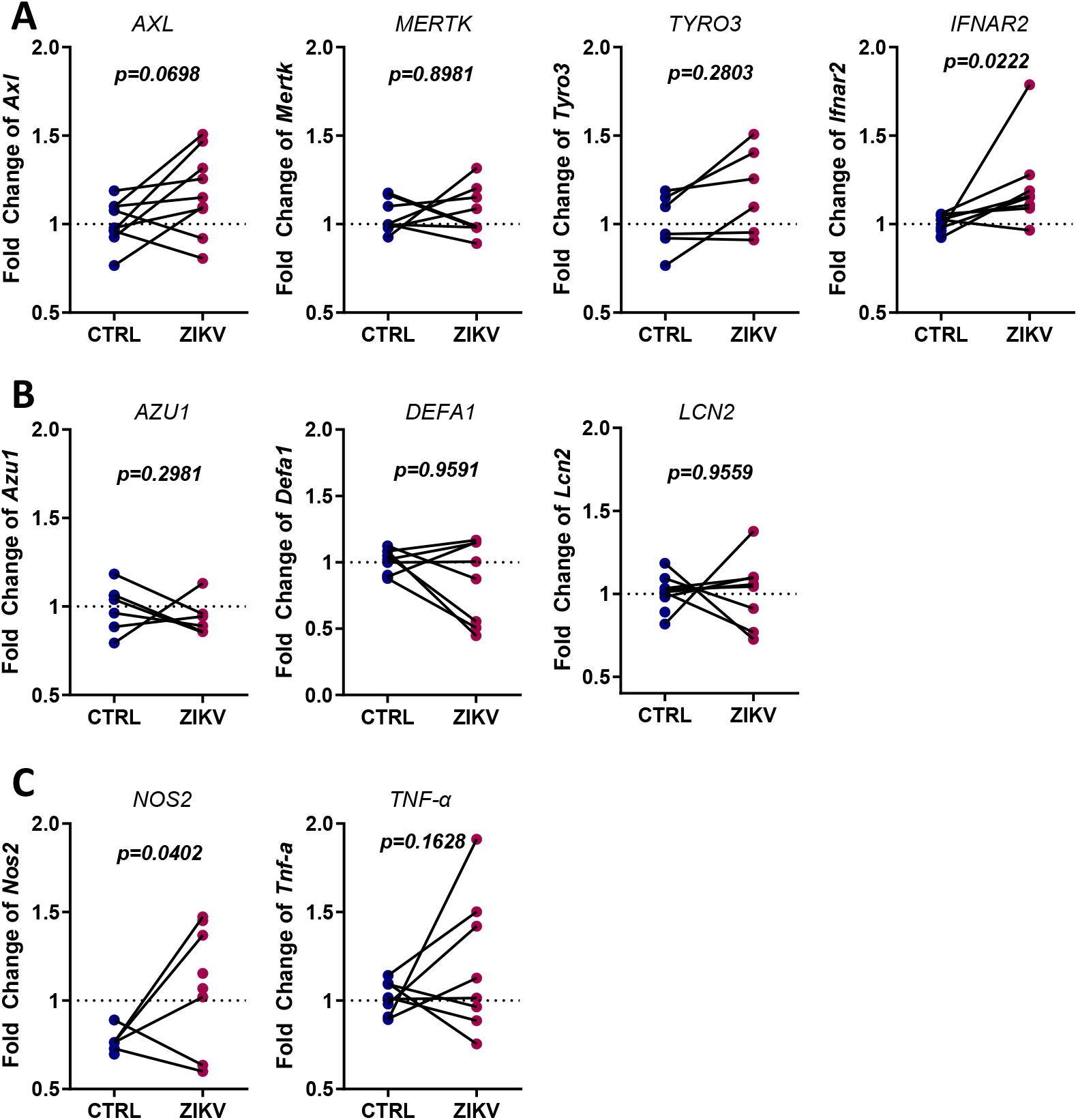
Gene expression profile in ZIKV-infected human neutrophils. Peripheral blood neutrophils from healthy subjects were infected in vitro with ZIKV (MOI 0.1) for 9 hours. Then, the total RNA was extracted, converted into cDNA and submitted to qPCR for A) TAM receptors (AXL, MERTK, TYRO3) and Interferon receptor IFNAR2, B) Neutrophil function (AZU1, DEFA1, LCN2) and C) Immune response (NOS2 and Tnfa). Data from 6-8 subjects with three replicates. Unpaired two-tailed t-test was used.

**Supplementary figure 6:**
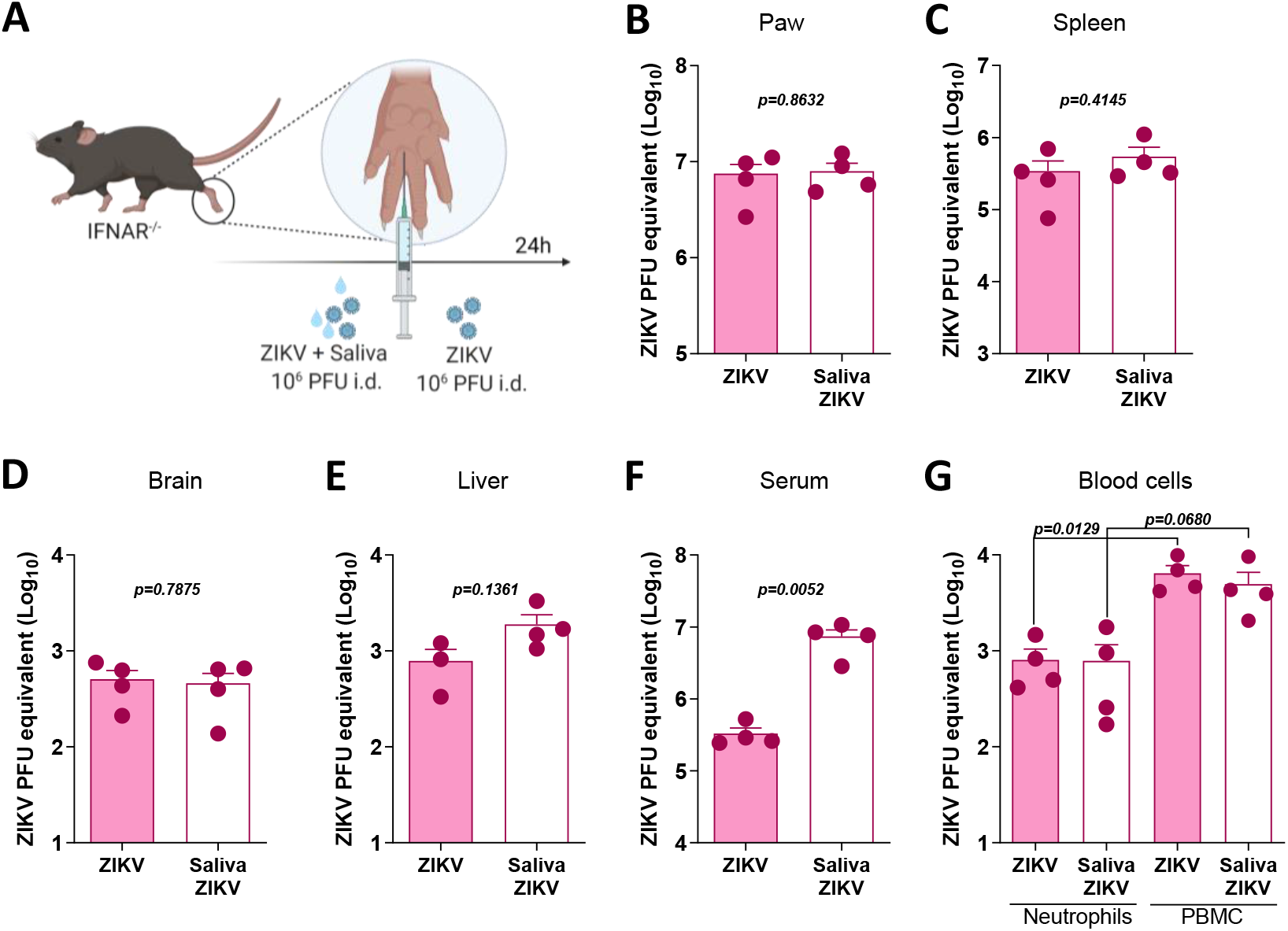
Aedes aegypti saliva induces higher viremia. A) Experimental design. C57BL/6 IFNAR-/- mice were infected with 106 PFU of ZIKV in the presence of 10ug of Aedes egypti saliva. Twenty-four hours later B) paw, C) spleen, D) brain, E) liver, F) serum and G) neutrophils and PBMC from blood were harvested submitted to viral RNA detection by RT-qPCR. Data representative of 3 experiments with 3 - 4 replicates. Unpaired two-tailed t-test was used from B to F. One-way ANOVA was used in G.

**Supplementary figure 7:**
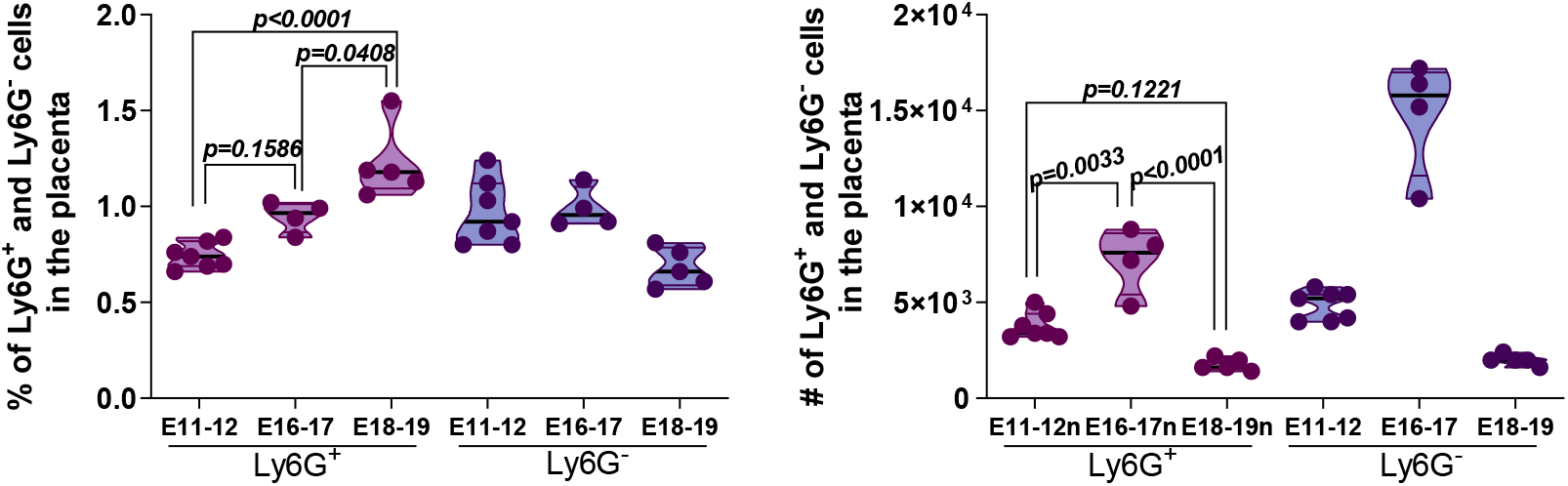
Frequency and number of neutrophils and macrophages in the placenta of IFNAR-/- mice. Placentas were harvested at the timepoints shown and chemically and mechanically dissociated for the measurement of the frequency and number of neutrophils (CD11b+Ly6G+) and macrophages (CD11b+Ly6G-) by flow cytometry. One-way ANOVA was used.

**Supplementary figure 8:**
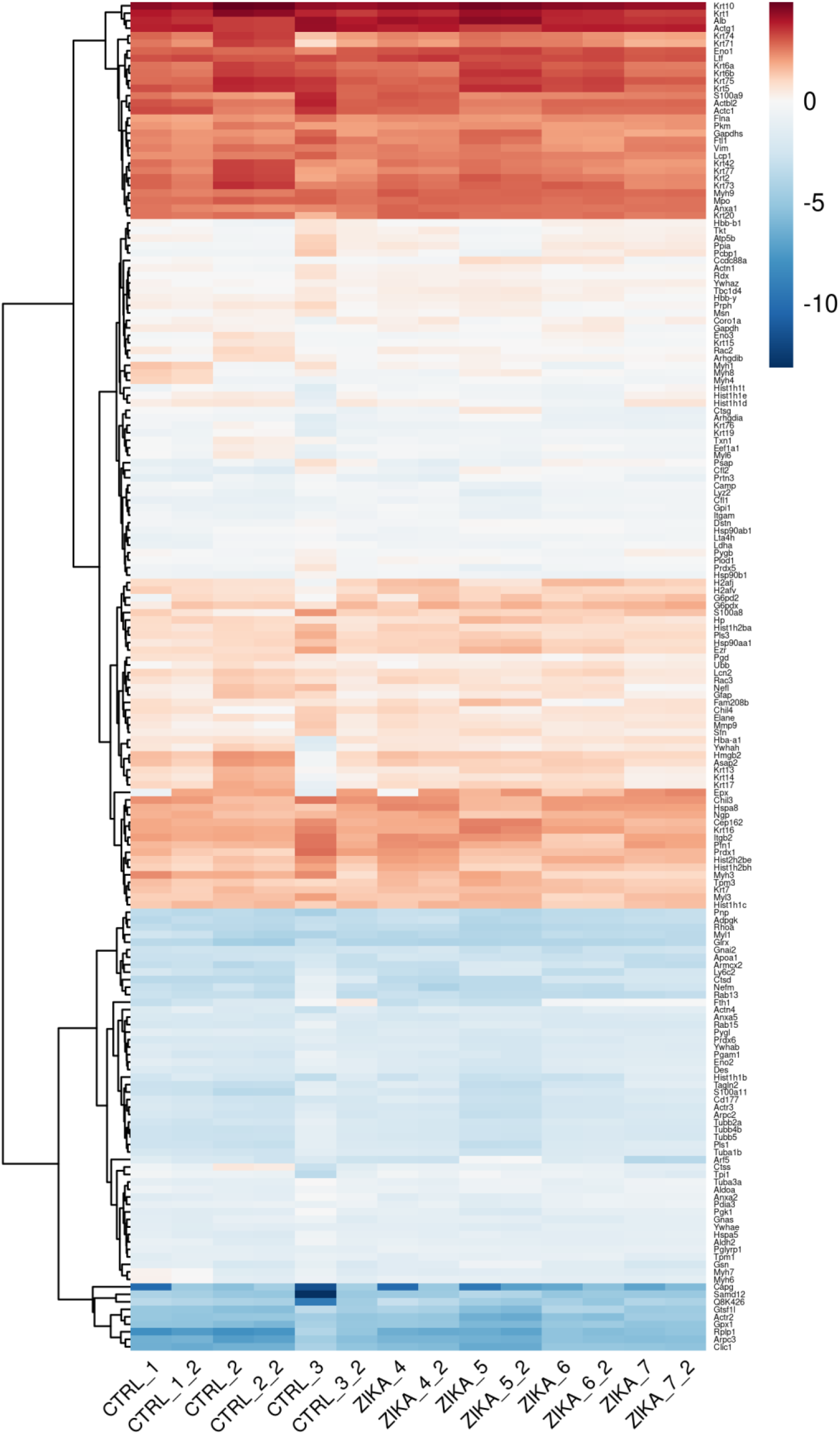
Coverage of proteomic analysis from placental neutrophils. Neutrophils isolated from placenta in figure 5 were submitted to protein extraction followed by proteomics analysis. Heatmap with 180 from 192 proteins detected in the proteomic analysis. All these proteins were not statistically different between control and ZIKV groups.

**Supplementary figure 9:**
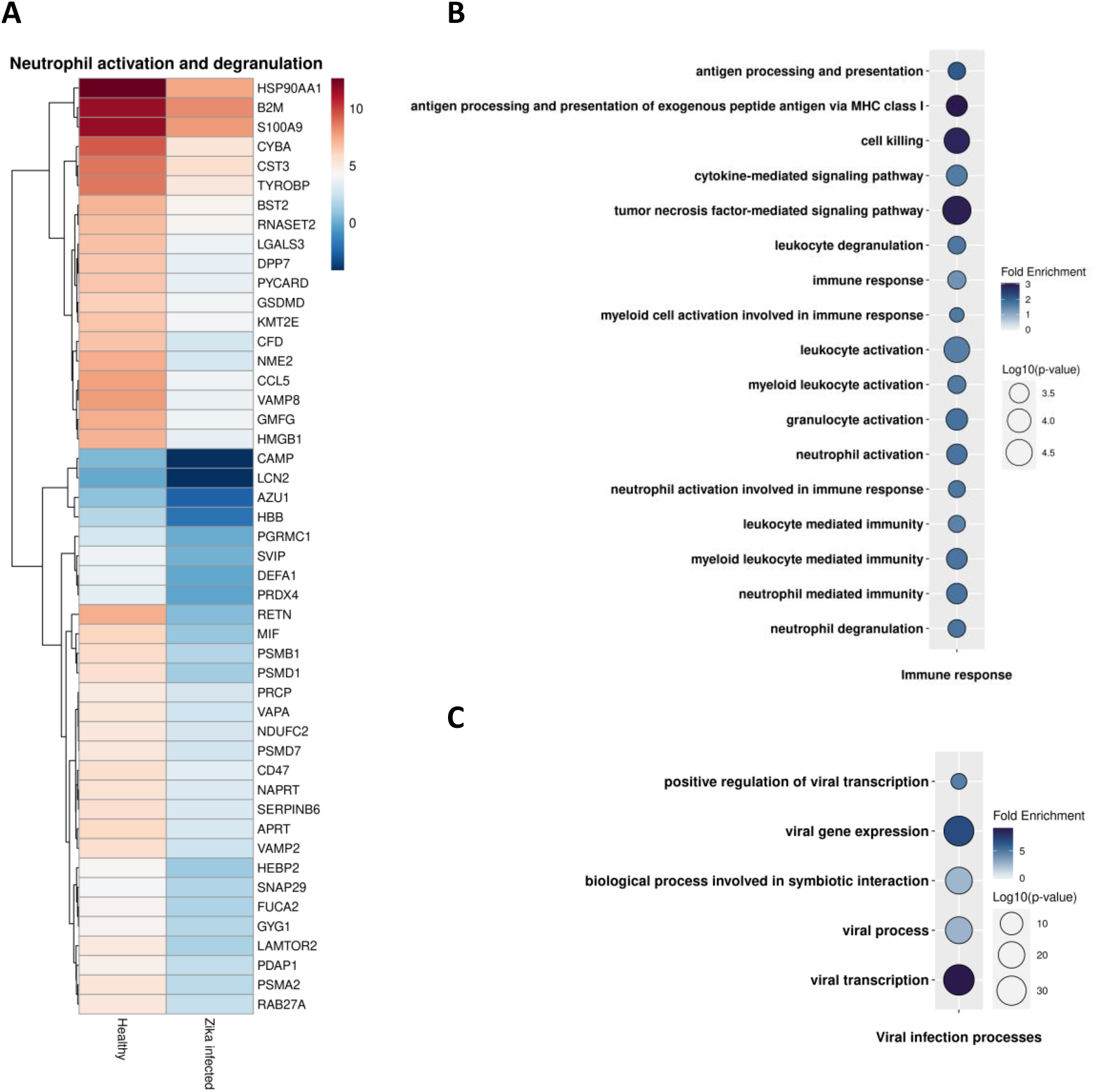
Genes downregulated in human ZIKV-infected placenta. Available RNAseq data set from Lum et al., 2019 of CD45+ cells from pre-term placenta of ZIKV-infected on the first trimester and health donor were analyzed. A) Genes involved in neutrophil activation and degranulation were downregulated in ZIKV-infected placenta. B-C) Gene ontology (GO) of downregulated genes in ZIKV-infected placenta showed enrichment of B) immune response pathways and C) viral infection processes.

## REFERENCES

1. R.-M. AJ, Zika: the new arbovirus threat for Latin America, J. Infect. Dev. Ctries. 9, 684–685 (2015).

2. F. NR, Q. J, C. IM, T. J, de J. JG, G. M, K. MUG, H. SC, B. A, da C. AC, F. LC, S. SP, W. CH, R. J, C. S, du P. L, V. MP, de O. WK, C. EH, C. GE, S. ACFS, V. LC, H. CM, S. JT, L. M, A. KG, G. ND, S. S, C. CY, M.-M. JE, G.-B. CR, A. CF, L.-X. LL, B. SA, C. AO, A. SF, F. CA, L. PS, N. BLS, M. HAO, S. IC, de Q. MG, de S. TR, B. JF, L. MR, P. GF, L. D, M. LC, D. R, F. RF, M. T, M. ET, J. T, W. GL, de L. MC, N. V, de C. EM, de L. MM, M. DL, N. JPM, L. AS, T.-M. TR, F. SN, M.-C. MC, M. FP, S. A, H. EC, R. A, B. T, N. MRT, S. EC, A. LCJ, L. NJ, P. OG, Establishment and cryptic transmission of Zika virus in Brazil and the Americas, Nature 546, 406–410 (2017).

3. L. C. Caires-Júnior, E. Goulart, U. S. Melo, B. H. S. Araujo, L. Alvizi, A. Soares-Schanoski, D. F. de Oliveira, G. S. Kobayashi, K. Griesi-Oliveira, C. M. Musso, M. S. Amaral, L. F. daSilva, R. M. Astray, S. F. Suárez-Patiño, D. C. Ventini, S. Gomes da Silva, G. L. Yamamoto, S. Ezquina, M. S. Naslavsky, K. A. Telles-Silva, K. Weinmann, V. van der Linden, H. van der Linden, J. R. M. de Oliveira, N. M. R. Arrais, A. Melo, T. Figueiredo, S. Santos, J. G. C. Meira, S. D. Passos, R. P. de Almeida, A. J. B. Bispo, E. A. Cavalheiro, J. Kalil, E. Cunha-Neto, H. Nakaya, R. Andreata-Santos, L. C. de Souza Ferreira, S. Verjovski-Almeida, P. L. Ho, M. R. Passos-Bueno, M. Zatz, Discordant congenital Zika syndrome twins show differential in vitro viral susceptibility of neural progenitor cells, Nat. Commun. 2018 91 9, 1–11 (2018).

4. F. R. Cugola, I. R. Fernandes, F. B. Russo, B. C. Freitas, J. L. M. Dias, K. P. Guimarães, C. Benazzato, N. Almeida, G. C. Pignatari, S. Romero, C. M. Polonio, I. Cunha, C. L. Freitas, W. N. Brandaõ, C. Rossato, D. G. Andrade, D. D. P. Faria, A. T. Garcez, C. A. Buchpigel, C. T. Braconi, E. Mendes, A. A. Sall, P. M. D. A. Zanotto, J. P. S. Peron, A. R. Muotri, P. C. B. B. Beltrao-Braga, The Brazilian Zika virus strain causes birth defects in experimental models, Nature 534 (2016), doi:10.1038/nature18296.

5. P. Brasil, G. A. Calvet, A. M. Siqueira, M. Wakimoto, P. C. de Sequeira, A. Nobre, M. de S. B. Quintana, M. C. L. de Mendona, O. Lupi, R. V. de Souza, C. Romero, H. Zogbi, C. da S. Bressan, S. S. Alves, R. Loureno-de-Oliveira, R. M. R. Nogueira, M. S. Carvalho, A. M. B. de Filippis, T. Jaenisch, Zika Virus Outbreak in Rio de Janeiro, Brazil: Clinical Characterization, Epidemiological and Virological Aspects, PLoS Negl. Trop. Dis. 10, 1–13 (2016).

6. J. J. Miner, B. Cao, J. Govero, K. K. Noguchi, I. U. Mysorekar, M. S. Diamond, J. J. Miner, B. Cao, J. Govero, A. M. Smith, E. Fernandez, O. H. Cabrera, C. Garber, M. Noll, R. S. Klein, K. K. Noguchi, I. U. Mysorekar, Zika Virus Infection during Pregnancy in Mice Causes Placental Damage and Fetal Demise Article Zika Virus Infection during Pregnancy in Mice Causes Placental Damage and Fetal Demise, Cell 165, 1081–1091 (2016).

7. C. M. Polonio, C. L. de Freitas, N. G. Zanluqui, J. P. S. Peron, Zika virus congenital syndrome: Experimental models and clinical aspects, J. Venom. Anim. Toxins Incl. Trop. Dis. 23, 1–9 (2017).

8. A. Ž. T. Mlakar J, Korva M, Tul N, Popović M, Poljšak-Prijatelj M, Mraz J, Kolenc M, Resman Rus K, Vesnaver Vipotnik T, Fabjan Vodušek V, Vizjak A, Pižem J, Petrovec M, V. F. Vodušek, A. Vizjak, D. Ph, J. Pižem, D. Ph, Zika Virus Associated with Microcephaly., N Engl J Med 374, 951–8 (2016).

9. V. Papayannopoulos, Neutrophil extracellular traps in immunity and disease Nat. Rev. Immunol. 18, 134–147 (2018).

10. C. H. Hiroki, J. E. Toller-Kawahisa, M. J. Fumagalli, D. F. Colon, L. T. M. Figueiredo, B. A. L. D. Fonseca, R. F. O. Franca, F. Q. Cunha, Neutrophil Extracellular Traps Effectively Control Acute Chikungunya Virus Infection, Front. Immunol. 10 (2020), doi:10.3389/fimmu.2019.03108.

11. F. P. Veras, M. C. Pontelli, C. M. Silva, J. E. Toller-Kawahisa, M. de Lima, D. C. Nascimento, A. H. Schneider, D. Caetité, L. A. Tavares, I. M. Paiva, R. Rosales, D. Colón, R. Martins, I. A. Castro, G. M. Almeida, M. I. F. Lopes, M. N. Benatti, L. P. Bonjorno, M. C. Giannini, R. Luppino-Assad, S. L. Almeida, F. Vilar, R. Santana, V. R. Bollela, M. Auxiliadora-Martins, M. Borges, C. H. Miranda, A. Pazin-Filho, L. L. P. da Silva, L. Cunha, D. S. Zamboni, F. Dal-Pizzol, L. O. Leiria, L. Siyuan, S. Batah, A. Fabro, T. Mauad, M. Dolhnikoff, A. Duarte-Neto, P. Saldiva, T. M. Cunha, J. C. Alves-Filho, E. Arruda, P. Louzada-Junior, R. D. Oliveira, F. Q. Cunha, SARS-CoV-2-triggered neutrophil extracellular traps mediate COVID-19 pathology, J. Exp. Med. 217 (2020), doi:10.1084/jem.20201129.

12. K. Ley, H. M. Hoffman, P. Kubes, M. A. Cassatella, A. Zychlinsky, C. C. Hedrick, S. D. Catz, Neutrophils: New insights and open questions, Sci. Immunol. 3, eaat4579 (2018).

13. R. Stillie, S. M. Farooq, J. R. Gordon, A. W. Stadnyk, The functional significance behind expressing two IL–8 receptor types on PMN, J. Leukoc. Biol. 86, 529–543 (2009).

14. V. Brinkmann, U. Reichard, C. Goosmann, B. Fauler, Y. Uhlemann, D. S. Weiss, Y. Weinrauch, A. Zychlinsky, Neutrophil Extracellular Traps Kill Bacteria, Science (80-.). 303, 1532–1535 (2004).

15. N. Branzk, V. Papayannopoulos, Molecular mechanisms regulating NETosis in infection and disease Semin. Immunopathol. 35, 513–530 (2013).

16. M. Pingen, S. R. Bryden, E. Pondeville, E. Schnettler, A. Kohl, A. Merits, J. K. Fazakerley, G. J. Graham, C. S. McKimmie, Host Inflammatory Response to Mosquito Bites Enhances the Severity of Arbovirus Infection, Immunity 44, 1455–1469 (2016).

17. M. M. B. Moreno-Altamirano, O. Rodríguez-Espinosa, O. Rojas-Espinosa, B. Pliego-Rivero, F. J. Sánchez-Garciá, Dengue Virus Serotype-2 Interferes with the Formation of Neutrophil Extracellular Traps, Intervirology 58, 250–259 (2015).

18. F. Bai, K. F. Kong, J. Dai, F. Qian, L. Zhang, C. R. Brown, E. Fikrig, R. R. Montgomery, A paradoxical role for neutrophils in the pathogenesis of West Nile virus, J. Infect. Dis. 202, 1804–1812 (2010).

19. Y. Zhao, M. Lu, L. T. Lau, J. Lu, Z. Gao, J. Liu, A. C. H. Yu, Q. Cao, J. Ye, M. A. McNutt, J. Gu, Neutrophils may be a vehicle for viral replication and dissemination in human h5n1 avian influenza, Clin. Infect. Dis. 47, 1575–1578 (2008).

20. Z. Zhang, T. Huang, F. Yu, X. Liu, C. Zhao, X. Chen, D. J. Kelvin, J. Gu, Infectious Progeny of 2009 A (H1N1) influenza virus replicated in and released from human neutrophils, Sci. Rep. 5, 1–11 (2015).

21. T. Saitoh, J. Komano, Y. Saitoh, T. Misawa, M. Takahama, T. Kozaki, T. Uehata, H. Iwasaki, H. Omori, S. Yamaoka, N. Yamamoto, S. Akira, Neutrophil Extracellular Traps Mediate a Host Defense Response to Human Immunodeficiency Virus-1, Cell Host Microbe 12, 109–116 (2012).

22. K. Lim, T. hyoun Kim, A. Trzeciak, A. M. Amitrano, E. C. Reilly, H. Prizant, D. J. Fowell, D. J. Topham, M. Kim, In situ neutrophil efferocytosis shapes T cell immunity to influenza infection, Nat. Immunol. 21, 1046–1057 (2020).

23. T. TM, C. SH, O. JE, L. RN, Neutrophil-mediated suppression of virus replication after herpes simplex virus type 1 infection of the murine cornea, J. Virol. 70, 898–904 (1996).

24. M. D. Tate, A. G. Brooks, P. C. Reading, J. D. Mintern, Neutrophils sustain effective CD8 T-cell responses in the respiratory tract following influenza infection, Immunol. Cell Biol. 90, 197–205 (2012).

25. H. E, G. B, L. W, W. W, O. Y, K. MJ, L. RI, H. BC, Human alpha- and beta-defensins block multiple steps in herpes simplex virus infection, J. Immunol. 177, 8658–8666 (2006).

26. C. B, de B. OJ, de J. R, A. AF, S. P. YS, L. R, van W. JB, B. RA, Neutrophil extracellular traps cause airway obstruction during respiratory syncytial virus disease, J. Pathol. 238, 401–411 (2016).

27. J. Heo, P. Dogra, T. J. Masi, E. A. Pitt, P. de Kruijf, M. J. Smit, T. E. Sparer, Novel Human Cytomegalovirus Viral Chemokines, vCXCL-1s, Display Functional Selectivity for Neutrophil Signaling and Function, J. Immunol. 195, 227–236 (2015).

28. A. Opasawatchai, P. Amornsupawat, N. Jiravejchakul, W. Chan-in, N. J. Spoerk, K. Manopwisedjaroen, P. Singhasivanon, T. Yingtaweesak, S. Suraamornkul, J. Mongkolsapaya, A. Sakuntabhai, P. Matangkasombut, F. Loison, Neutrophil activation and early features of net formation are associated with dengue virus infection in human, Front. Immunol. 10, 3007 (2019).

29. M. Pingen, S. R. Bryden, E. Pondeville, E. Schnettler, A. Kohl, A. Merits, J. K. Fazakerley, G. J. Graham, C. S. McKimmie, Host Inflammatory Response to Mosquito Bites Enhances the Severity of Arbovirus Infection, Immunity 44, 1455–1469 (2016).

30. F. R. Cugola, I. R. Fernandes, F. B. Russo, B. C. Freitas, J. L. M. Dias, K. P. Guimarães, C. Benazzato, N. Almeida, G. C. Pignatari, S. Romero, C. M. Polonio, I. Cunha, C. L. Freitas, W. N. Brandaõ, C. Rossato, D. G. Andrade, D. D. P. Faria, A. T. Garcez, C. A. Buchpigel, C. T. Braconi, E. Mendes, A. A. Sall, P. M. D. A. Zanotto, J. P. S. Peron, A. R. Muotri, P. C. B. B. Beltrao-Braga, The Brazilian Zika virus strain causes birth defects in experimental models, Nature 534, 267–271 (2016).

31. E. Hatanaka, A. C. Levada-Pires, T. C. Pithon-Curi, R. Curi, Systematic study on ROS production induced by oleic, linoleic, and γ-linolenic acids in human and rat neutrophils, Free Radic. Biol. Med. 41, 1124–1132 (2006).

32. V. Brinkmann, B. Laube, U. A. Abed, C. Goosmann, A. Zychlinsky, Neutrophil extracellular traps: How to generate and visualize them, J. Vis. Exp. 1724 (2010).

33. M. Arenas-Hernandez, E. N. Sanchez-Rodriguez, T. N. Mial, S. A. Robertson, N. Gomez-Lopez, Isolation of leukocytes from the murine tissues at the maternal-fetal interface, J. Vis. Exp. 2015, 52866 (2015).

34. R. S. Lanciotti, O. L. Kosoy, J. J. Laven, J. O. Velez, A. J. Lambert, A. J. Johnson, S. M. Stanfield, M. R. Duffy, Genetic and serologic properties of Zika virus associated with an epidemic, Yap State, Micronesia, 2007, Emerg. Infect. Dis. 14, 1232–1239 (2008).

35. W. JR, O. P, M. M, High recovery FASP applied to the proteomic analysis of microdissected formalin fixed paraffin embedded cancer tissues retrieves known colon cancer markers, J. Proteome Res. 10, 3040–3049 (2011).

36. S. JC, G. MV, L. GZ, V. JP, G. SJ, Absolute quantification of proteins by LCMSE: a virtue of parallel MS acquisition, Mol. Cell. Proteomics 5, 144–156 (2006).

37. V. JP, L. JI, A. JM, Analysis and quantification of diagnostic serum markers and protein signatures for Gaucher disease, Mol. Cell. Proteomics 6, 755–766 (2007).

38. B. AM, L. M, U. B, Trimmomatic: a flexible trimmer for Illumina sequence data, Bioinformatics 30, 2114–2120 (2014).

39. S. JT, D. R, Efficient de novo assembly of large genomes using compressed data structures, Genome Res. 22, 549–556 (2012).

40. D. Kim, J. M. Paggi, C. Park, C. Bennett, S. L. Salzberg, Graph-based genome alignment and genotyping with HISAT2 and HISAT-genotype, Nat. Biotechnol. 2019 378 37, 907–915 (2019).

41. B. G, H. KE, K. EJ, D. C. B, F. GA, G. GR, Simulation-based comprehensive benchmarking of RNA-seq aligners, Nat. Methods 14, 135–139 (2017).

42. A. S, P. PT, H. W, HTSeq--a Python framework to work with high-throughput sequencing data, Bioinformatics 31, 166–169 (2015).

43. M. I. Love, W. Huber, S. Anders, Moderated estimation of fold change and dispersion for RNA-seq data with DESeq2, Genome Biol. 2014 1512 15, 1–21 (2014).

44. H. Mi, A. Muruganujan, X. Huang, D. Ebert, C. Mills, X. Guo, P. D. Thomas, Protocol Update for large-scale genome and gene function analysis with the PANTHER classification system (v.14.0), Nat. Protoc. 2019 143 14, 703–721 (2019).

45. D. Michlmayr, P. Andrade, K. Gonzalez, A. Balmaseda, E. Harris, CD14+CD16+ monocytes are the main target of Zika virus infection in peripheral blood mononuclear cells in a paediatric study in Nicaragua, Nat. Microbiol. 2, 1462–1470 (2017).

46. M. O. Henrique, L. S. Neto, J. B. Assis, M. S. Barros, M. L. Capurro, A. P. Lepique, D. M. Fonseca, A. Sá-Nunes, Evaluation of inflammatory skin infiltrate following Aedes aegypti bites in sensitized and non-sensitized mice reveals saliva-dependent and immune-dependent phenotypes, Immunology 158, 47–59 (2019).

47. F. M. Lum, V. Narang, S. Hue, J. Chen, N. McGovern, R. Rajarethinam, J. J. L. Tan, S. N. Amrun, Y. H. Chan, C. Y. P. Lee, T. K. Chua, W. X. Yee, N. K. W. Yeo, T. C. Tan, X. Liu, S. Haldenby, Y. sin Leo, F. Ginhoux, J. K. Y. Chan, J. Hiscox, C. Y. Chong, L. F. P. Ng, Immunological observations and transcriptomic analysis of trimester-specific full-term placentas from three Zika virus-infected women, Clin. Transl. Immunol. 8, 1–15 (2019).

48. J. L. Silva-Filho, L. G. de Oliveira, L. Monteiro, P. L. Parise, N. G. Zanluqui, C. M. Polonio, C. L. de Freitas, D. A. Toledo-Teixeira, W. M. de Souza, N. Bittencourt, M. R. Amorim, J. Forato, S. P. Muraro, G. F. de Souza, M. C. Martini, K. Bispo-dos-Santos, A. Vieira, C. C. Judice, G. M. Pastore, E. Amaral, R. Passini Junior, H. M. B. P. Mayer-Milanez, C. C. Ribeiro-do-Valle, R. Calil, J. Renato Bennini Junior, G. J. Lajos, A. Altemani, M. T. Nolasco da Silva, A. Carolina Coan, M. Francisca Colella-Santos, A. P. B. von Zuben, M. A. R. Vinolo, C. W. Arns, R. R. Catharino, M. L. Costa, R. N. Angerami, A. R. R. Freitas, M. R. Resende, M. T. Garcia, M. Luiza Moretti, L. Renia, L. F. P. Ng, C. V. Rothlin, F. T. M. Costa, J. P. S. Peron, J. L. Proença-Modena, Gas6 drives Zika virus-induced neurological complications in humans and congenital syndrome in immunocompetent mice, Brain. Behav. Immun. (2021), doi:10.1016/j.bbi.2021.08.008.

49. P. G. Quie, J. G. White, B. Holmes, R. A. Good, In Vitro Bactericidal Capacity of Human Polymorphonuclear Leukocytes: Diminished Activity in Chronic Granulomatous Disease of Childhood, J. Clin. Invest. 46, 668 (1967).

50. W. K, Z. C, D. DC, Severe congenital neutropenia, Semin. Hematol. 43, 189–195 (2006).

51. M. Pingen, S. R. Bryden, E. Pondeville, E. Schnettler, A. Kohl, A. Merits, J. K. Fazakerley, G. J. Graham, C. S. McKimmie, Host Inflammatory Response to Mosquito Bites Enhances the Severity of Arbovirus Infection, Immunity 44, 1455–1469 (2016).

52. M. M, C. SS, O. GG, K. WV, R. G, F. CL, S. DL, P. WD, K. DB, S. AL, Activation of triggering receptor expressed on myeloid cells-1 on human neutrophils by marburg and ebola viruses, J. Virol. 80, 7235–7244 (2006).

53. A. K. Hastings, R. Uraki, H. Gaitsch, K. Dhaliwal, S. Stanley, H. Sproch, E. Williamson, T. MacNeil, A. Marin-Lopez, J. Hwang, Y. Wang, J. R. Grover, E. Fikrig, Aedes aegypti NeSt1 Protein Enhances Zika Virus Pathogenesis by Activating Neutrophils, J. Virol. 93 (2019), doi:10.1128/jvi.00395-19.

54. P. Mistry, S. Nakabo, L. O’Neil, R. R. Goel, K. Jiang, C. Carmona-Rivera, S. Gupta, D. W. Chan, P. M. Carlucci, X. Wang, F. Naz, Z. Manna, A. Dey, N. N. Mehta, S. Hasni, S. Dell’Orso, G. Gutierrez-Cruz, H. W. Sun, M. J. Kaplan, Transcriptomic, epigenetic, and functional analyses implicate neutrophil diversity in the pathogenesis of systemic lupus erythematosus, Proc. Natl. Acad. Sci. U. S. A. 116, 25222–25228 (2019).

55. C. A. Prada-Medina, J. P. S. Peron, H. I. Nakaya, Immature neutrophil signature associated with the sexual dimorphism of systemic juvenile idiopathic arthritis, J. Leukoc. Biol. 108, 1319–1327 (2020).

56. J. Mestas, C. C. W. Hughes, Of Mice and Not Men: Differences between Mouse and Human Immunology, J. Immunol. 172, 2731–2738 (2004).

57. G. Boivin, J. Faget, P. B. Ancey, A. Gkasti, J. Mussard, C. Engblom, C. Pfirschke, C. Contat, J. Pascual, J. Vazquez, N. Bendriss-Vermare, C. Caux, M. C. Vozenin, M. J. Pittet, M. Gunzer, E. Meylan, Durable and controlled depletion of neutrophils in mice, Nat. Commun. 11, 1–9 (2020).

58. D. S, S. X, L. S, W. L, C. V, R. Y, Interleukin-10-mediated inhibition of free radical generation in macrophages, Am. J. Physiol. Lung Cell. Mol. Physiol. 280 (2001), doi:10.1152/AJPLUNG.2001.280.6.L1196.

59. E. Camacho-Zavala, C. Santacruz-Tinoco, E. Muñoz, R. Chacón-Salinas, M. I. Salazar-Sanchez, C. Grajales, J. González-Ibarra, V. H. Borja-Aburto, T. Jaenisch, C. R. Gonzalez-Bonilla, Pregnant Women Infected with Zika Virus Show Higher Viral Load and Immunoregulatory Cytokines Profile with CXCL10 Increase, Viruses 2021, Vol. 13, Page 80 13, 80 (2021).

60. Z. J, S. O, S. S, B. A, B. L, L. SJ, Immunophenotyping and activation status of maternal peripheral blood leukocytes during pregnancy and labour, both term and preterm, J. Cell. Mol. Med. 21, 2386–2402 (2017).

61. L. R, S. S, A. R, P. R, Granulocyte superoxide anion production and regulation by plasma factors in normal and preeclamptic pregnancy, J. Reprod. Immunol. 89, 199–206 (2011).

62. S. Nadkarni, J. Smith, A. N. Sferruzzi-Perri, A. Ledwozyw, M. Kishore, R. Haas, C. Mauro, D. J. Williams, S. H. P. Farsky, F. M. Marelli-Berg, M. Perretti, Neutrophils induce proangiogenic T cells with a regulatory phenotype in pregnancy, Proc. Natl. Acad. Sci. 113, E8415–E8424 (2016).

63. R. AZ, Y. W, H. DA, R. CA, S. DA, Placental Pathology of Zika Virus: Viral Infection of the Placenta Induces Villous Stromal Macrophage (Hofbauer Cell) Proliferation and Hyperplasia, Arch. Pathol. Lab. Med. 141, 43–48 (2017).

64. K. A. Jurado, M. K. Simoni, Z. Tang, R. Uraki, J. Hwang, S. Householder, M. Wu, B. D. Lindenbach, V. M. Abrahams, S. Guller, E. Fikrig, Zika virus productively infects primary human placenta-specific macrophages, JCI Insight 1, 88461 (2016).

65. G. R. Santos, C. A. L. Pinto, R. C. S. Prudente, S. S. Witkin, A. S. Arandes, L. C. Rodrigues, M. Zatz, E. Massad, S. D. Passos, Differences in Placental Histology Between Zika Virus–infected Teenagers and Older Women, Int. J. Gynecol. Pathol. Publish Ahead of Print, 1–8 (2021).

66. A. T. L. Horvath, K. M. Milano, E. Fikrig, T. Rakib, L. J. Yockey, A. Millet, A. Iwasaki, H. J. Kliman, C. B. Coyne, S. Weatherbee, K. A. Jurado, N. Arora, Y. Kong, K. Hastings, Type I interferons instigate fetal demise after Zika virus infection, Sci. Immunol. 3, eaao1680 (2018).

67. S. K, H. GD, Emerging role for AS160/TBC1D4 and TBC1D1 in the regulation of GLUT4 traffic, Am. J. Physiol. Endocrinol. Metab. 295 (2008), doi:10.1152/AJPENDO.90331.2008.

68. Z. QL, J. ZY, H. J, C. A, H. GN, L. J, C. MP, Akt substrate TBC1D1 regulates GLUT1 expression through the mTOR pathway in 3T3-L1 adipocytes, Biochem. J. 411, 647–655 (2008).

69. H. AFA, M. P, C. AS, A. M, E. F, M. R, J. P, WNK1 phosphorylation sites in TBC1D1 and TBC1D4 modulate cell surface expression of GLUT1, Arch. Biochem. Biophys. 679 (2020), doi:10.1016/J.ABB.2019.108223.

70. K. Maher, B. J. Kokelj, M. Butinar, G. Mikhaylov, M. Manček-Keber, V. Stoka, O. Vasiljeva, B. Turk, S. A. Grigoryev, N. Kopitar-Jerala, A Role for Stefin B (Cystatin B) in Inflammation and Endotoxemia *, J. Biol. Chem. 289, 31736–31750 (2014).

71. M. Butinar, M. T. Prebanda, J. Rajković, B. Jerič, V. Stoka, C. Peters, T. Reinheckel, A. Krüger, V. Turk, B. Turk, O. Vasiljeva, Stefin B deficiency reduces tumor growth via sensitization of tumor cells to oxidative stress in a breast cancer model, Oncogene 2014 3326 33, 3392–3400 (2013).

72. Z. Q, P. QZ, Z. AL, H. H, Z. JJ, T. Y, H. WM, L. M, W. DS, C. MY, M. G, X. JC, Annexin A3 upregulates the infiltrated neutrophil-lymphocyte ratio to remodel the immune microenvironment in hepatocellular carcinoma, Int. Immunopharmacol. 89 (2020), doi:10.1016/J.INTIMP.2020.107139.

73. M. SE, M. RO, The annexins, Genome Biol. 5 (2004), doi:10.1186/GB-2004-5-4-219.

74. S. He, X. Li, R. Li, L. Fang, L. Sun, Y. Wang, M. Wu, Annexin A2 Modulates ROS and Impacts Inflammatory Response via IL-17 Signaling in Polymicrobial Sepsis Mice, PLOS Pathog. 12, e1005743 (2016).

75. L. Yang, P. Lu, X. Yang, K. Li, S. Qu, Annexin A3, a Calcium-Dependent Phospholipid-Binding Protein: Implication in Cancer, Front. Mol. Biosci. 0, 700 (2021).

76. L. C. V, M.-P. I, Annexin 3 is associated with cytoplasmic granules in neutrophils and monocytes and translocates to the plasma membrane in activated cells, Biochem. J. 303 ( Pt 2, 481–487 (1994).

77. E. G. Kallas, L. G. F. A. B. D’Elia Zanella, C. H. V. Moreira, R. Buccheri, G. B. F. Diniz, A. C. P. Castiñeiras, P. R. Costa, J. Z. C. Dias, M. P. Marmorato, A. T. W. Song, A. Maestri, I. C. Borges, D. Joelsons, N. B. Cerqueira, N. C. Santiago e Souza, I. Morales Claro, E. C. Sabino, J. E. Levi, V. I. Avelino-Silva, Y. L. Ho, Predictors of mortality in patients with yellow fever: an observational cohort study, Lancet Infect. Dis. 19, 750–758 (2019).

78. K. S. A. Khabar, F. Al-Zoghaibi, M. N. Al-Ahdal, T. Murayama, M. Dhalla, N. Mukaida, M. Taha, S. T. Al-Sedairy, Y. Siddiqui, G. Kessie, K. Matsushima, The α Chemokine, Interleukin 8, Inhibits the Antiviral Action of Interferon α, J. Exp. Med. 186, 1077 (1997).

79. J. Wagoner, M. Austin, J. Green, T. Imaizumi, A. Casola, A. Brasier, K. S. A. Khabar, T. Wakita, M. Gale, S. J. Polyak, Regulation of CXCL-8 (Interleukin-8) Induction by Double-Stranded RNA Signaling Pathways during Hepatitis C Virus Infection, J. Virol. 81, 309–318 (2007).

80. T. Pollicino, L. Bellinghieri, A. Restuccia, G. Raffa, C. Musolino, A. Alibrandi, D. Teti, G. Raimondo, Hepatitis B virus (HBV) induces the expression of interleukin-8 that in turn reduces HBV sensitivity to interferon-alpha, Virology 444, 317–328 (2013).

81. M. N, Pathophysiological roles of interleukin-8/CXCL8 in pulmonary diseases, Am. J. Physiol. Lung Cell. Mol. Physiol. 284 (2003), doi:10.1152/AJPLUNG.00233.2002.

